# CLONAL LYMPHOCYTE EXPANSIONS AND JAK-STAT PATHWAY MUTATIONS DEFINE A PATHOGENIC CONTINUUM DRIVING RESISTANCE TO GLUTEN-FREE DIET IN CELIAC DISEASE

**DOI:** 10.1101/2025.08.27.672584

**Authors:** Georgia Malamut, Nicolas Guégan, Francesco Carbone, Cécile Masson, Marine Luka, Sascha Cording, Christine Bole, Mathurin Fumery, Guillaume Bouguen, Vered Abitbol, Romain Coriat, Benoît Terris, Barbara Burroni, Frédéric Beuvon, Ludovic Lhermitte, Chantal Brouzes, Henri Duboc, Isabelle Nion-Larmurier, Mathieu Uzzan, David Sibon, Vahid Asnafi, Olivier Hermine, Mickaël Ménager, Nadine Cerf-Bensussan, Anaïs Levescot

## Abstract

**Background&Aims:** Despite recent advances, refractory celiac disease (RCD) poses challenging questions. In type 2 RCD (RCD2), the lack of response to the gluten-free diet is attributed to an intestinal intraepithelial lymphoma carrying driver JAK1 or STAT3 mutations. However, it remains unclear whether these can be safely targeted for therapy. In RCD1, pathogenic insights are still lacking.

**Methods:** Duodenal biopsies and peripheral blood mononuclear cells (PBMCs) from patients with RCD1, RCD2, active CeD, CeD in remission, and controls were analyzed. Lymphocyte populations were characterized using single-cell transcriptomic, genomic, and TCR repertoire profiling. Functional and exome sequencing analyses were performed on patient-derived RCD2 cell lines exposed to JAK inhibitors.

**Results:** We show that clonal malignant RCD2 lymphocytes exhibit interpatient similarities but substantial intratumoral heterogeneity, and provide *in vitro* evidence that JAK inhibitors can select drug-resistant tumor cells, arguing against their use as monotherapy. In RCD1, we identified clonal T-cell expansions harboring mutations that enhance the JAK-STAT pathway. The detection of both RCD2 and a CD4⁺ lymphoproliferation in a patient initially diagnosed with RCD1 further illustrates the diversity of lymphoproliferative outcomes in celiac disease.

**Conclusions:** These findings suggest that RCD subtypes may share underlying mechanisms driven by clonal evolution and JAK-STAT dysregulation. They also highlight the potential limitations of JAK inhibitor monotherapy and the importance of molecularly informed therapeutic strategies.

**What You Need to Know:** *BACKGROUND AND CONTEXT:* Refractory celiac disease (RCD) can lead to intestinal lymphoma, but the biological processes driving immune cell transformation and therapy resistance remain incompletely understood.

*NEW FINDINGS:* Single-cell analyses reveal clonal evolution, JAK-STAT pathway dysregulation, and shared molecular features between RCD subtypes, with implications for disease progression and treatment response.

*LIMITATIONS:* Sample size and reliance on *ex vivo* models limit generalizability; further *in vivo* validation of resistance mechanisms is needed to confirm therapeutic implications.

*CLINICAL RESEARCH RELEVANCE:* These findings suggest that RCD subtypes may share underlying mechanisms driven by clonal evolution and JAK-STAT dysregulation. They also highlight the potential limitations of JAK inhibitor monotherapy and the importance of molecularly informed therapeutic strategies.

*BASIC RESEARCH RELEVANCE:* This work uncovers mechanisms of immune cell transformation in chronic intestinal inflammation and provides insight into how tumor heterogeneity and somatic mutations drive disease progression and therapeutic resistance.

**Lay summary:** This study reveals shared mechanisms in refractory celiac disease subtypes driven by clonal evolution and JAK-STAT activation, highlighting limits of JAK inhibitor monotherapy and the need for personalized treatments.

## INTRODUCTION

Celiac disease (CeD), which affects 1% of the global population, exemplifies how interactions between environmental and genetic factors can trigger chronic autoimmune-like inflammation and ultimately lead to lymphomagenesis ^1–3^. CeD is driven by the activation of CD4⁺ T cells by gluten peptides presented by HLA-DQ2 or HLA-DQ8, the main genetic predisposing factors^2–4^. The gluten-specific CD4^+^ T cells produce cytokines, notably IFNγ, IL-2, IL-21 which, alongside IL-15 produced by myeloid and epithelial cells in the celiac intestine, promote the destruction of the gut epithelium by cytotoxic CD8^+^ intraepithelial T lymphocytes (IELs)^3,5,6^. CeD is generally controlled by a strict gluten-free diet (GFD). However, some patients develop resistance to the GFD^7–12^. A well-established cause of refractoriness, commonly referred to as type 2 refractory CeD (RCD2), is a low-grade lymphoma characterized by the clonal expansion of innate-like intestinal IELs, which typically lack surface expression of T-cell markers but express intracellular CD3 and contain clonal T cell receptor DNA rearrangements^12,13^. The prognosis is complicated by malnutrition and frequent progression to enteropathy-associated T-cell lymphomas (EATL)^7,8,10,12,14^. The identification of somatic gain-of-function (GOF) mutations in JAK1 and STAT3 in nearly all RCD2 and EATL cases has revealed a key mechanism linking gluten-driven chronic inflammation to lymphomagenesis by promoting the outgrowth of malignant IELs in response to cytokines produced in the celiac gut^12,13,15,16^. A second group of patients refractory to GFD are classified as RCD1^7,8,11,12^. Traditionally considered as a non-clonal and less severe disease than RCD2, its mechanism remains largely elusive (reviewed in^12^). However, recent findings of clonal T-cell expansions with lymphoma-associated mutations^17^ suggest potential for clonal evolution, warranting further investigation.

Both RCD1 and RCD2 generally respond to corticoids. Yet, RCD2 patients may develop resistance and progress to EATL^7,8,11,12,18^. The high prevalence of JAK1 and STAT3 GOF mutations has prompted the use of JAK inhibitors. Although these drugs improved histological lesions, they failed to reduce RCD2 IEL numbers^19–21^, suggesting that they may alleviate inflammation whithout eliminating malignant cells. Genomic analysis indicates that RCD2 malignant cells accumulate multiple oncogenic mutations beyond JAK1/STAT3 ^15^. Such a high mutational burden is a plausible source of intra-tumor heterogeneity (ITH), a major challenge for targeted therapies^22^.

Here, single cell approaches were used to further characterize the pathogenic intestinal lymphocytes involved in the loss of response to GFD in CeD patients and to assess how ITH and evolutionary dynamics may favor resistance to targeted treatment. In RCD2, clonal innate-like malignant IELs showed substantial transcriptomic and genomic ITH. Moreover, exposure of an RCD2 cell lines to a JAK inhibitor led to the recurrent selection of drug-resistant subclones, carrying mutations that were absent or barely detectable in the parental population, highlighting a potential risk of clonal selection under targeted therapy in RCD2. In RCD1, the identification of clonal T-cell expansions carrying somatic mutations that enhance the JAK-STAT pathway and the possible progression of RCD1 into RCD2 support the hypothesis of a mechanistic overlap or a continuum between the two types of RCD.

## MATERIAL AND METHODS

### Patients

Patients with RCD2 (n=12), RCD1 (n=4), active celiac disease (aCeD, n=1), CeD in remission on a gluten-free diet (GFD, n=5), and non-CeD controls with normal duodenal histology (n=4) were enrolled after informed consent, as described in the supplementary material and summarized in **Table S1**. All patients with CeD (refractory or not) had positive anti-TG2 antibodies at diagnosis and carried the HLA-DQ2 or -DQ8 haplotype when tested. At the time of sample collection, anti-TG2 antibodies were negative in all but the aCeD patient.

### Cell preparation from blood and biopsies for single cell transcriptomic studies

Cryopreserved biopsies were thawed and washed in pre-warmed (37°C) RPMI medium (Invitrogen) containing 5% fetal bovine serum (FBS) (Invitrogen). After transfer in gentleMACS™ C Tubes, biopsies were digested using the Multi Tissue Dissociation Kit 1 and the GentleMACS dissociator (all from Miltenyi Biotec) according to the Miltenyi Program 37C_h_TDK_1 and manufacturer’s instructions. After further tissue dissociation by several flushes through a 20mL syringe with a 18 gauge needle, the cell suspension was filtered through a 100µm cell strainer. Cells were centrifuged and labelled with anti-human CD45-PerCP/Cy5.5 (Clone HI30, SONY Biotechnology Inc) and the fixable aqua LIVE/DEAD (ThermoFisher Scientific) and live CD45^+^ cells were sorted using a SONY MA900 Multi-Application Cell Sorter (SONY Biotechnology Inc). Cryopreserved PBMCs were thawed in warmed RPMI medium with 10% FBS.

### Single-cell RNA sequencing

Sorted CD45^+^ intestinal cells (10^5^-2×10^5^) and 2×10^5^ PBMCs were labelled with TotalSeqTM Human Universal Cocktail V1.0 for 30 minutes at 4°C according to manufacturer’s instructions. After controlling by trypan blue exclusion that viability of PBMCs and CD45^+^ intestinal cells was >90%, these cells were encapsulated into droplets using the 10x Genomics Chromium Single Cell platform to create 5’ single-cell cDNA libraries for gene expression, cell surface protein, and TCR sequencing. The libraries were prepared according to the manufacturer’s protocol and sequenced on an Illumina NovaSeq platform. Since initial studies had shown over 90% viability and minimal transcriptome variability in CD45^+^ cells from fresh and cryopreserved biopsies, all experiments were performed using cryopreserved material (**Figure S1**).

### scRNA-seq Data Processing

Samples were collected and processed over several years. Sequencing reads were demultiplexed and aligned to the human reference transcriptome (GRCh38 and GRCh38-2020-A) using the CellRanger pipeline (v4, v5, and v7). Unfiltered raw UMI counts from CellRanger were imported into Seurat (v4.0.4) for quality control, data integration, and subsequent analyses. Cells with fewer than 500 features or a mitochondrial content exceeding 20% were excluded to remove empty beads, apoptotic cells, and potential doublets.

For each sample, data were normalized and scaled using the log normalization method. The top 2,000 highly variable genes were identified with the FindVariableFeatures function. Batch effects across samples were corrected using Seurat’s FindIntegrationAnchors function, and integration was performed using canonical correlation analysis (CCA). Dimensionality reduction was conducted on the integrated dataset using principal component analysis (PCA), computed from the first 20 PCs. The k-nearest neighbor graph was constructed using the FindNeighbors function, and cell clusters were identified using FindClusters at multiple resolutions (0.8 to 2.0). For downstream analysis and visualization, the maximum resolution (r = 1.2) was selected, aligning best with expected cell populations.

### Whole Exome Sequencing

Genomic DNA was purified using the QIAMP DNA mini Kit in accordance with the manufacturer’s instructions (Qiagen). Extracted DNA was quality-controlled and assayed using Xpose instrument (Trinean). Exome capture was performed using the Twist Bioscience kit. Twist exome libraries were prepared from 30ng to 3 µg of fragmented genomic DNA using an ultrasonicator (Covaris), in accordance with the manufacturer’s recommendations. Barcoded exome libraries were pooled and sequenced with a Novaseq6000 system (Illumina), generating paired-end reads. After demultiplexing, sequences were aligned to the human genome reference (NCBI build 37, version hg19) using Illumina DRAGEN software. After demultiplexing, variant detection was performed with the DRAGEN software suite. The average depth of coverage of exome libraries was over 120X, with over 96-99% of target exonic bases covered by at least 15 independent reads and over 93-98% by at least 30 independent reads (98-99% at 15X and 94-97% at 30X). All variants were annotated and filtered with PolyWeb, an annotation software developed in-house.

For other methods, please see supplemental data

## RESULTS

### Single-cell transcriptomic mapping and lineage tracing of RCD2 Tumor Cells

To establish the transcriptomic landscape of RCD lymphocytes, single-cell RNA sequencing (scRNA-seq) was performed on CD45^+^ cells sorted from cryopreserved duodenal biopsies and peripheral blood mononuclear cells (PBMCs) of patients with RCD2 (n=12), RCD1 (n=4), active CeD (n=1), CeD in remission on a GFD (n=5), and control individuals with normal duodenal histology (n= 4) (**Figure 1A-B**, **Table S1**). Live cells were encapsulated in droplets to generate single-cell libraries for gene expression, cell-surface protein, and αβ-TCR sequencing (**Figure 1B**). A total of 217,749 cells from 38 biopsies passed quality control. Unsupervised clustering using canonical immune markers identified 19 immune cell types, including T cells, B cells, NK cells, and mast cells^23^ (**Figure 1C**). Since RCD2 tumor cells were rare in PBMCs (**Table S1**), analyses focused on biopsy-derived cells.

**Figure 1.**
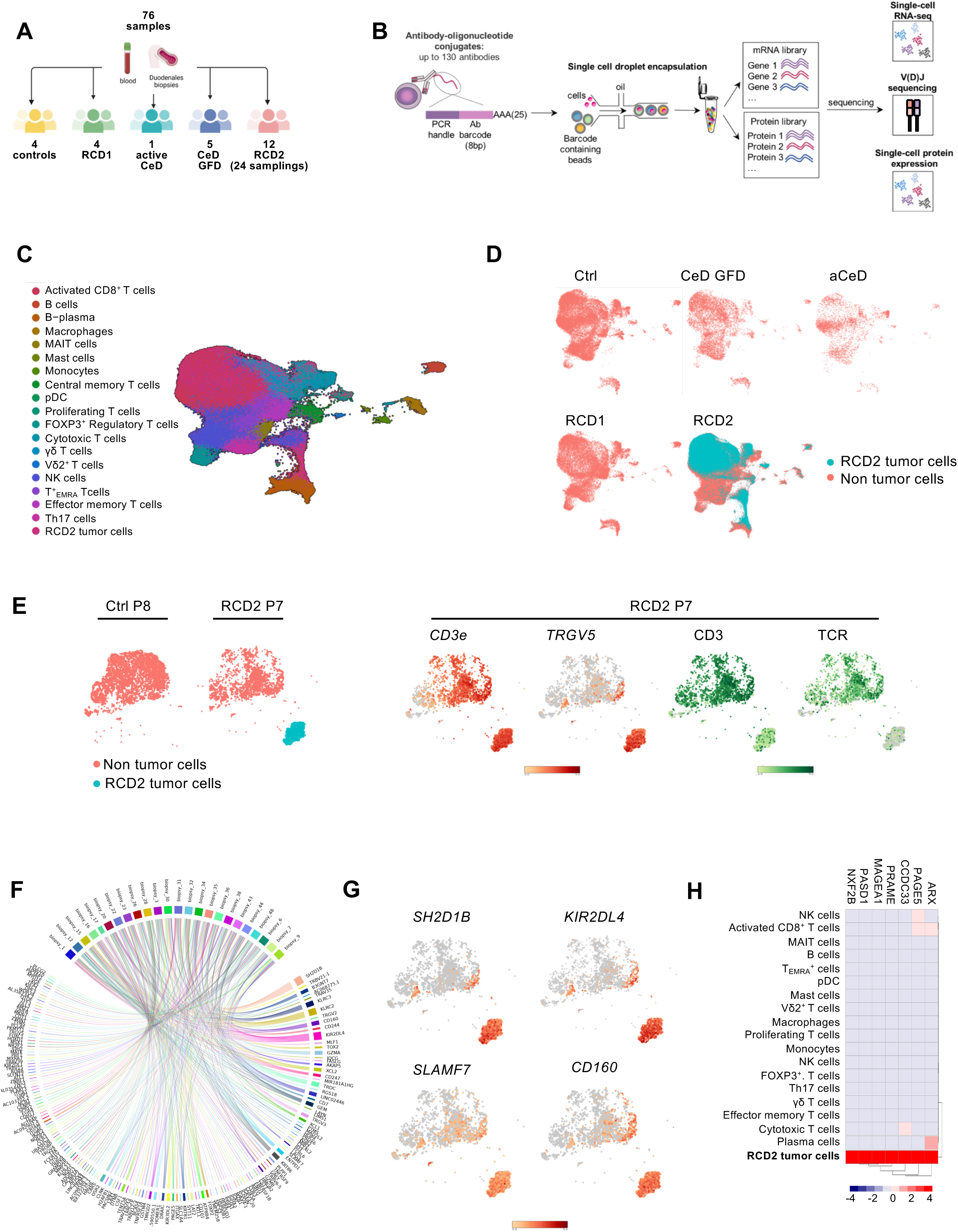
Single-cell transcriptomic mapping and lineage tracing of RCD2 tumor cells. **A**. Overview of analyzed samples. **B.** Workflow combining scRNA-seq, TCR αβ sequencing, and surface protein profiling. **C-D.** UMAP of 217,749 CD45^+^ cells from 38 samples across RCD2, RCD1, and controls and annotation of malignant RCD2 IELs (in blue). **E.** Annotation of malignant RCD2 cells in indicated individual biopsies based on gene and protein expression (orange-to-red and green gradients, respectively). **F.** Circos plot of top 20 genes overexpressed in RCD2 tumor cells. **G.** UMAP highlighting expression of NK-associated transcripts. **H.** Heatmap showing cancer-testis gene expression in RCD2 versus non-tumor CD45⁺ cells.

Tumor cells identification was challenging when RCD2 biopsies where pooled (**Figure 1D**). However, individual biopsies analysis revealed clusters exclusively present in RCD2 patients **(Figure 1E**). In these clusters, presence of RCD2 tumor cells was confirmed by absent surface CD3E and TCR expression, contrasting with the presence of transcripts encoding CD3E and a single TCRG variable chain (**Figure 1E**). Tumor-to-immune cell ratios varied markedly across biopsies, even within the same patient over time, reflecting tumor burden and/or varying degrees of immune infiltration (**Figures 1D** and **S2**). Comparison of normal duodenal immune cells confirmed that RCD2 tumor cells express hallmark genes of both ILC1/NK cells, and CD8^+^ T-IELs, and are more closely related to activated CD8^+^ T-IELs than to NK cells, consistent with prior bulk RNA analysis^13^ (**Figures 1E** and **S3**).

In line with previous work suggesting occasional clonal TCR α or β chains rearrangements in RCD2^13,24^, full-length V(D)J sequencing identified in-frame TCR α and/or β transcripts in 6 patients (**Figure S4**). In 4 cases, RCD2 cells expressed a single in-frame α or β chain transcript (**Figure S4**). Unexpectedly, in 2 other cases, tumor cells expressing a single TCRG variable transcript showed multiple in-frame α and/or β chains (**Figure S4**), suggesting ongoing rearrangements. However, the absence of detectable RAG recombinases transcripts prevented confirmation of recombination activity.

Some transcripts, notably *SH2D1B* and *KIR2DL4*, were consistently overexpressed in RCD2 cells compared to other immune subsets, providing robust markers even when surface CD3 labelling was suboptimal (**Figure 1F-G**). *SH2D1B* encodes the intracellular adaptor EAT-2, which enables NK cell cytotoxicity via the transmembrane protein SLAMF7/CRACC/CS1^25^, whose transcripts were also often upregulated in RCD2 tumor cells (**Figure 1F**-**G**). *KIR2DL4* encodes an immunoglobulin-like transmembrane receptor that is expressed by NK cells and a subset of CD8^+^ T cells and can inhibit cytotoxicity ^26^. Both transcripts were also markedly increased in one atypical RCD2 case (P28), in which tumor cells expressed surface CD3 and TCRγδ, a phenotype reported in rare patients^12,15,16^. These cells displayed a transcriptomic profile largely similar to other RCD2 samples, along with uniformly strong expression of *TRGV2* transcripts, consistent with the clonal *TCRG* rearrangement identified in isolated IELs (**Figure S5**, **Table S1**). Additionally, RCD2 cells expressed cancer-testis genes such as *MAGEA1*, *PAGE5*^27^, and *PASD1*, which may promote STAT3-dependent signaling and tumorogenesis^28^ (**Figure 1H**). Cancer-testis gene expression, typically restricted to the male germline, can be reactivated in tumors. Expression of cancer-testis genes varied across patients but was absent in normal cells, highlighting their biomarker potential^27^.

### Transcriptional evidence of inter-tumor similarity but high intratumor heterogeneity in RCD2

To assess transcriptomic heterogeneity within and across RCD2 patients, we refined clustering of 45,248 tumor cells pooled from 24 biopsies of 12 RCD2 patients (1 to 4 samples per patient, collected approximately one year apart; see **Table S1**). This analysis identified 24 transcriptional subclusters (**Figures 2A–B** and **S6**) and, strikingly, revealed strong inter-patient similarity, with few patient-specific clusters, consistent with a shared cellular origin and transformation mechanism^12,13^. In contrast, ITH was high, increasing with tumor burden, clinical severity (**figure 2C**) and tumor progression. In patient P7, the number of tumor clusters increased from 17 to 21 between the initial (P7) and the second (P7.2) assessment conducted one year later before extra-intestinal dissemination and death (**Figures 2B** and **S6**). In patient P1, ITH decreased alongside increasing budesonide dosage, from 17 clusters at baseline (3 mg/day), to 10 clusters (6 mg/day), and 11 clusters (9 mg/day) at subsequent assessments. However, cluster 1 expanded over time, increasing from 33% to 67.3% of tumor cells, coinciding with progression to EATL (**Figures 2B** and **S6**). In patients P12 and P22, initial assessments were conducted six months after chemotherapy and autologous stem cell transplantation for EATL. Follow-up biopsies showed an increase in ITH, particularly at the third time point, one year prior EATL relapse and death: from 19 to 21 clusters in P12, and from 15 to 24 clusters in P22 (**Figures 2B** and **S6**). Similarly, in P3, the number of tumor clusters rose from 19 to 25 shortly before EATL diagnosis, with new subpopulations bearing chromosomal gains (P3.2) (**Figures 2B**,**D** and **S6**).

**Figure 2.**
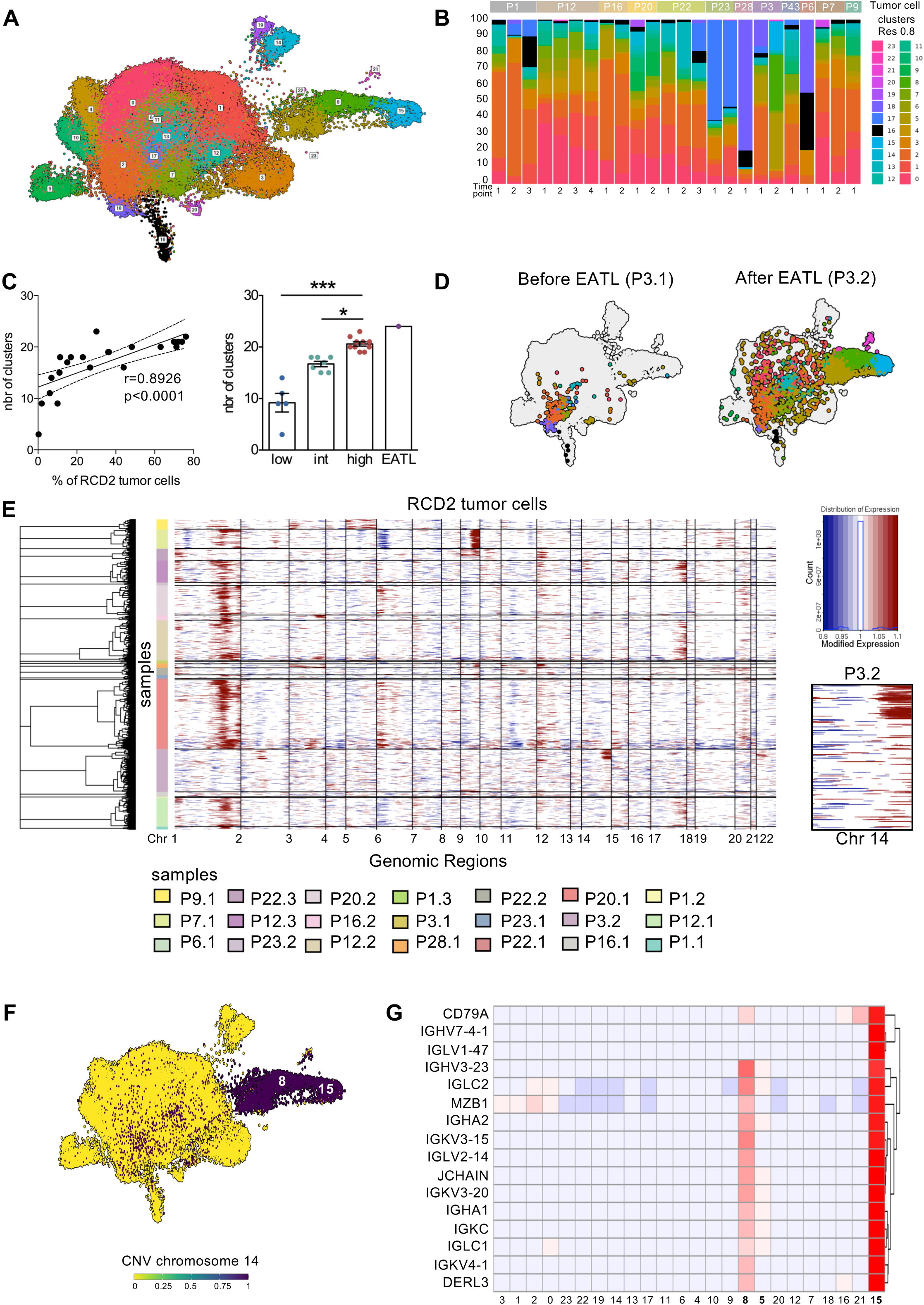
Gene expression, tumor cell clustering and trajectory analysis in RCD2. **A.** UMAP of 45,248 tumor cells isolated from 24 samples (12 RCD2 patients, 1 to 4 samples per patient) **B.** Proportions of cells per cluster. Cluster 16, enriched in proliferating cells, is shown in black **C.** Correlation between the number of tumor clusters and the percentages of tumor cells in each biopsy (left panel). Bar plot showing the number of tumor clusters in duodenal biopsies from RCD2 patients with low, intermediate (int), and high tumor burden or with EATL (right panel). **D.** UMAPs visualization of RCD2 tumor cluster from patient P3 before (P3.1) and after EATL diagnosis (P3.2). **E.** CNV profiles inferred from scRNA-seq data obtained by comparing individual RCD2 tumor and non tumor cells (see **Figure S7**). The y-axis represents individual cells, and the x-axis corresponds to genomic position. *Inset:* zoom-in on chromosome 14 from patient P3 after EATL diagnosis. **F.** UMAP showing 14q gain localized within clusters 5, 8, and 15. **G.** Heatmap displaying genes upreguled in clusters 8 and 15 from patient P3 after EATL diagnosis. which harbor the 14q gain.

### Genomic evidence of inter- and intra-tumor heterogeneity in RCD2

The above findings highlighted the high degree of transcriptional ITH in RCD2 cells. However, such heterogeneity may arise from environmental cues or reflect underlying genomic diversity. To gain further insight into genomic ITH in RCD2, we used InferCNV^29^ to infer copy number variations (CNVs) from single-cell data. Consistent with previous findings in cell lines^15,30^, a recurrent 1q21–q32 gain was detected in primary RCD2 tumor cells from 6 patients (P1, P7, P12, P16, P20, P23) (**Figures 2E** and **S7**). The 1q gain is a well-characterized event in various cancers and may occur early during transformation^31,32^. Among the numerous genes encoded by the 1q21-32 region, several may contribute to lymphomagenesis, including *MDM4* (a negative regulator of P53), *BCL9* (a positive regulator of β-catenin-dependent transcription) and *MCL1* (an anti-apoptotic protein)^31,32^. Additional CNVs varied across patients, indicating inter-tumor genomic heterogeneity (**Figures 2E** and **S7**). Notably, in patient P3, and consistent with a possible link between genomic ITH and tumor progression, a 14q gain was observed in 3 tumor clusters that emerged (clusters 8 and 15) or expanded (cluster 5) following EATL diagnosis (**Figure 2E-F**). These clusters showed upregulation of IGHV7-4-1, IGHV3-23, IGHA1, and IGHA2, encoded by the immunoglobulin heavy chain (IGH) locus on 14q32 and normally restricted to B cells, suggesting transcriptional dysregulation and lineage deviation linked to the 14q gain (**Figures 2G** and **S8**).

### Tracking the evolutionary path of RCD2 cells from a proliferative cell cluster

In line with previous reports, most RCD2 cells lacked a proliferative signature^12,15,33^. However, pathway enrichment analysis revealed one cluster (cluster 16) overexpressing cell cycle genes *MKI67* and *STMN1* (**Figure 3A-B**). This cluster was detected in all biopsies, even in low-burden samples (**Figures 3B** and **S6**). For instance, patient P6, initially diagnosed with 50% RCD2 IELs (**Table S1**) showed a strong response to budesonide, with tumor cells representing only 0.3% of CD45^+^ cells at the time of initial sampling. However, 35% of those tumor cells belonged to cluster 16 (**Figure S6**), suggesting that this cluster may be enriched in cells with proliferative potential and could contribute to RCD2 persistence. Supporting this hypothesis, pseudotime analysis revealed a branched architecture originating from this cluster, with terminal states including clusters observed after EATL onset (**Figures 3B** and **S6**). While pseudotime analysis maps gene espression dynamics, it does not infer directionality or real-time kinetics. To address this question, we applied RNA velocity analysis using scVelo alogorithm, which estimates transcriptional dynamics based on the relative ratio of intronic (unspliced, immature) and exonic (spliced, mature) reads from scRNA-seq data^34^. This revealed a directional flow originating from the proliferative cluster 16 (**Figure 3C**). This pattern was further validated by partition-based graph abstraction (PAGA) analysis^35^, which identified this cluster as the likely origin point from which other tumor cell states emerge (**Figure 3C**). These results support the idea that the proliferative cluster contains tumor stem-like cells capable of seeding other RCD2 subpopulations.

**Figure 3:**
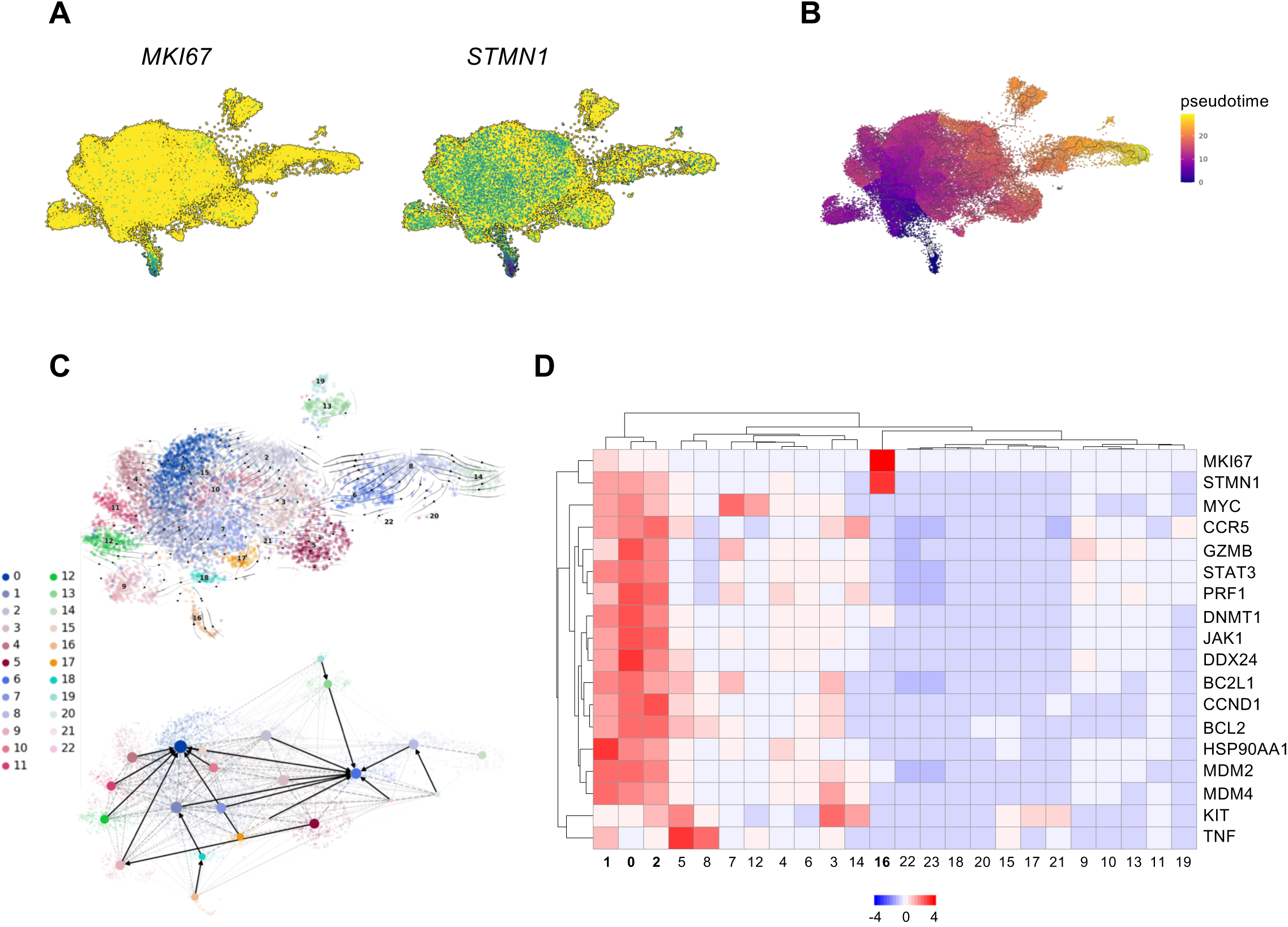
Evolutionary trajectory of RCD2 tumor cells originating from a proliferative cluster. **A.** Expression of proliferation markers *MKI67* and *STMN1* across tumor cells. **B.** Pseudotime projection of tumor cell differentiation on UMAP. **C.** *Top:* RNA velocity vectors inferred from the dynamical model, shown as streamlines over the UMAP-based embedding of RCD2 tumor cells. *Bottom:* Topology-preserving partition-based graph abstraction (PAGA) graph overlaid on the UMAP, visualizing potential cell-type transitions. Nodes represent tumor cell groups, and edge weights reflect the degree of connectivity. **D.** Heatmap of selected transcripts across tumor clusters. Two-way ANOVA with multiple comparisons. All results are represented as median + standard deviation (*p ≤ 0.05 to ****p ≤ 0.0001).

Interestingly, this proliferative cluster expressed low levels of *JAK1* and *STAT3*, contrasting with clusters 0, 1 and 2, which exhibited enhanced JAK/STAT signaling by pathway analysis (**Figure S9**) and strong expression of multiple STAT3 target genes, including *GZMB* and *PRF1* implicated in RCD2-mediated epithelial cytotoxicity^36^ (**Figure 3D**). These observations are consistent with clinical findings showing that JAK inhibitors can improve histologic lesions without reducing tumor burden in RCD2 patients^19–21^, possibly due to their selective activity against the inflammatory STAT3-high component of the tumor, while sparing the proliferative, stem-like cells.

### Selection of somatic mutations during acquisition of *in vitro* resistance to JAK inhibitors by RCD2 cells

To further investigate genomic ITH in RCD2, primary RCD2 tumor cells were sorted from frozen PBMCs collected at 3 different time points from a previously described patient (P3 in^15^) and analyzed by single-cell DNA sequencing with the Tapestri platform^37^ and a custom panel of 210 amplicons covering known RCD2-associated genomic alterations^15^ (SNVs and small indels) (**Table S2**). This analysis revealed 4 clones with increasing numbers of mutations, confirming genomic ITH and suggesting hierarchical clonal phylogeny (**Figure 4A**). The putative founder clone (clone 1) carried JAK1 p.G1097D and TNFAIP3 p.I36Gfs*61 mutations (**Figure 4A**), while later clones showed additional alterations in PIK3CG p.T887A (clone 2), STAT5B p.E433K (Clone 3), and STAT3 p.E616G (Clone 4). Clonal heterogeneity was also evident in both blood- and intestine-derived cell lines from the same patient with at least 4 different clones (**Figure 4B**).

**Figure 4:**
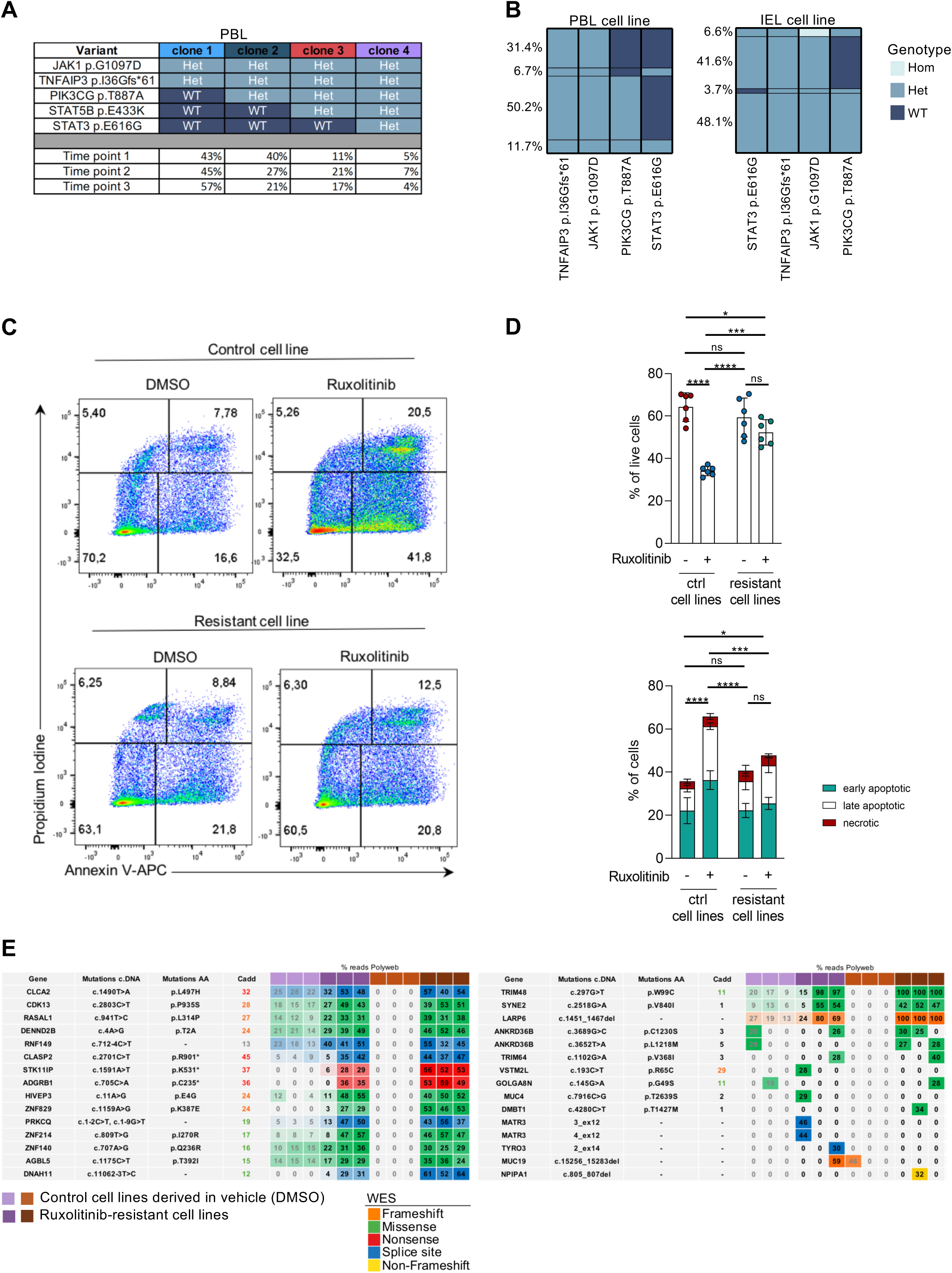
Evidence of subclonal genomic heterogeneity in RCD2 and *in vitro* selection of ruxolitinib-resistant cell lines. **A-B.** Targeted scDNA-seq of RCD2 cells purified from the peripheral blood of one RCD2 patient (P3 in^15^), collected at 3 distinct time points (**A**), and from two cell lines derived from the peripheral blood (PBL cell line) and intraepithelial lymphocyte (IEL) cell line of the same patient (**B**). **C-D.** Flow cytometry analysis comparing apoptosis in ruxolitinib-resistant versus control cell lines after 48h in the presence of IL-15 (20 ng/mL) and ruxolitinib (400nM) or DMSO. Significant differences were determined using a Kruskal–Wallis test followed by Dunn’s post hoc test for multiple comparisons (*p ≤ 0.05 to ****p ≤ 0.0001). **E.** Comparison of somatic mutations differentially expressed between 6 ruxolitinib-resistant cell lines and 6 control lines cultured with DMSO only. Mutations are color-coded by variant type, with numbers indicating variant allele frequency. Combined Annotation Dependent Depletion (Cadd) scores indicate predicted pathogenicity.

To assess how such heterogeneity might influence therapeutic response, the intestinal RCD2 cell line was treated over repeated cycles with the JAK inhibitor ruxolitinib, or maintained in vehicle-only conditions (DMSO), generating 6 resistant and 6 control lines. Upon re-exposure to ruxolitinib, resistant lines showed reduced apoptosis, indicating acquired resistance (**Figure 4C–D**). Whole-exome sequencing (WES) of resistant versus control lines identified 30 mutations (mean: 20 per resistant cell line) (**Figures 4E** and **Table S3**). Several mutations, including *CDK13* c.2803C>T and *RASAL1* c.941T>C, were recurrent across the resistant lines, while undectable or present at low read counts in control lines (**Figures 4E** and **Table S3**). These data support the hypothesis that ruxolitinib treatment selectively expands rare tumor subpopulations harboring co-selected mutations that may contribute to drug resistance, either individually or collectively. The RASAL1, CDK13 and CLASP2 variants are strong candidates due to their high pathogenicity scores and prior implication in tumor suppressor pathways^38–40^, along with a variant in ADGRB1, a known regulator of p53^41^ (**Figure S10**).

### Co-occurrence of RCD2 and CD4⁺ T-cell lymphoproliferation complicating RCD1

Patient P43 was initially enrolled as a case of RCD1. CeD was diagnosed in 1995, and RCD1 in 2005, at a time when duodenal biopsies showed ≤2% CD103^+^sCD3^-^ IELs by flow cytometry and no evidence of clonal TCRG rearrangement. However, in 2023, single-cell transcriptomic analysis revealed a clonal CD45⁺ population typical of RCD2, with an in-frame TCRβ transcript (*TRBV15/TRBJ1-2*) but no corresponding α-chain (**Figure 5A-D**), no surface CD3 by CITEseq but abundant *CD3E* and *SH2D1B* transcripts (**Figure 5D**). These cells accounted for 13.8% of live CD45^+^ cells in the duodenal biopsy and 1.7% in PBMCs. RCD2 was further confirmed by flow cytometry, which showed 37% of CD103⁺sCD3⁻iCD3⁺ IELs, and by bulk DNA analysis of IELs revealing a clonal TCRG rearrangement (**Table S1**). Unexpectedly, single-cell analysis also identified a clonal CD4⁺ T-cell lymphoproliferation accounting for 25% of circulating T cells, which was not detected in the duodenal compartment. These clonal cells exhibited low surface CD8 expression but high levels of CD56 and CD57 (**Figure 5A**, **C**, **E**), expressed in-frame TCR α (TRAV8-2/TRAJ10 ) and β (TRBV20-1/TRBJ2-3) transcripts, resulting in surface expression of the TCR Vβ2 chain (**Figure 5C**).

**Figure 5.**
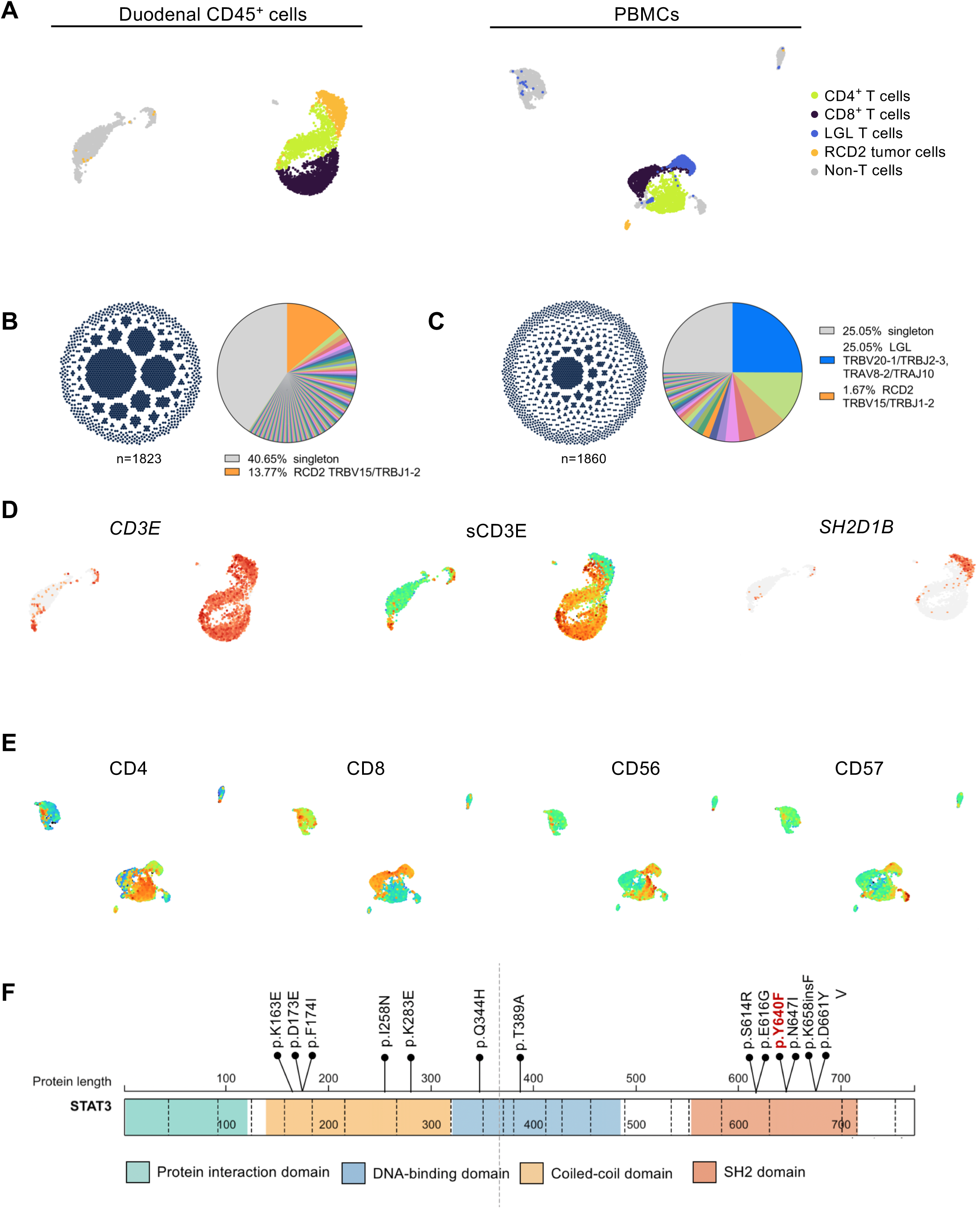
RCD2 and CD4^+^ lymphoproliferation complicating RCD1. **A.** UMAP of duodenal CD45^+^ cells (left panel) and PBMCs (right panel) from patient P43 showing the expanded CD4^+^ T cells clonotype (blue) and the RCD2 cells (orange). **B-C** Clonotype distribution ain duodenal CD45^+^ cells (**B**) and PBMCs (**C**) from patient P43 based on TCR α and β sequencing. Dots represent single cells grouped within clonotypes. Larger clonotypes appear centrally, smaller ones at the periphery. n indicates the number of cells with assigned paired TCRα and β sequences. **D.** Expression of *CD3E* and *SH2D1B* transcripts and surface CD3E protein in duodenal CD45^+^ cells from P43. **E.** Expression of surface protein CD4, CD8, CD56, and CD57 in PBMCs from P43. **F.** Localisation of the STAT3 variant identified in RCD2 cells of P43 (in red) compared to other STAT3 mutations previously reported in RCD2 patients^15^.

WES was performed on sorted CD103⁺sCD3⁻ RCD2 cells (from blood and biopsies), Vβ2⁺ CD4⁺ T cells from the lymphoproliferative population (from blood), and control peripheral Vβ2⁻ T cells. RCD2 cells harbored approximately 650 somatic mutations, consistent with the high mutational burden previously reported in RCD2 cell lines^15^ (**Table S4**), and carried both in blood and biopsy a heterozygous GOF STAT3 p.Y640F mutation (**Figure 5F** and **Table S4**), frequently found in LGL leukemia^42,43^. This mutation was absent in the Vβ2⁺ cells, excluding a shared clonal origin. No canonical LGL mutations were detected in Vβ2⁺ cells^43^, but 41 de novo mutations were identified, including a heterozygous missense mutation in ATM (c.7422A>C, CADD score 25) (**Table S5**). ATM is frequently mutated in lymphoid malignancies^44^. Moreover, biased usage of the TRBV20-1 gene segment has been reported across several major T-cell lymphoma subtypes^45,46^, supporting the notion that this population may represent a transformed clone. These findings demonstrate that two independent lymphoproliferative disorders, RCD2 and a CD4⁺ T-cell lymphoproliferation, emerged in patient P43 over the 18 years following initial RCD1diagnosis.

### Clonal expansion of duodenal cytotoxic CD8^+^ T cells harboring somatic SOCS1 and SOCS3 mutations in RCD1

Single-cell TCR analysis confirmed that oligoclonal T cell expansions are common in intestinal biopsies from controls^47^ (**Figure S11**). Expanded clones clustering with other T cells were also detected in 3 of 4 RCD1 patients (**Figure S11**). However, in patient P10, a large duodenal expansion of CD8⁺ TCRαβ⁺ T cells formed a distinct transcriptional cluster in UMAP space, suggesting a divergent functional state (**Figure 6A-F**). Full-length TCRαβ sequencing revealed a single clonotype expressing a Vβ1⁺ chain (*TRBV9/TRBJ2-7)* paired with two α chains (*TRAV-38-1/TRAJ53* and *TRAV36/DV7/TRAJ4)*. These clonal CD8^+^ T cells represented ∼21% of duodenal T cells (**Figure 6A-C**) and 10% of peripheral T cells (**Figure 6B-D**) which strongly expressed the CD103 protein, indicating an intestinal origin (**Figure 6F).** In both compartments, these cells displayed a distinct transcriptional profile compared to other CD8⁺ T cells from the same patient, characterized by elevated expression of cytotoxic (*GZMB*, *PRF1*), Th1 (*IFNG*), and NK-associated transcripts (*KLRC1*, *HCST, SH2D1B*, *CD160*) (**Figures 6G-H** and **S12**). Circulating clonal CD8^+^ T cells also upregulated *CCR9,* a chemokine receptor promoting homing to the small intestine, suggesting possible recirculation between intestinal sites(**Figure 6H**). A subset of these cells, especially in the blood, expressed *MKI67*, indicating proliferative activity. (**Figure 6 E-G**).

**Figure 6.**
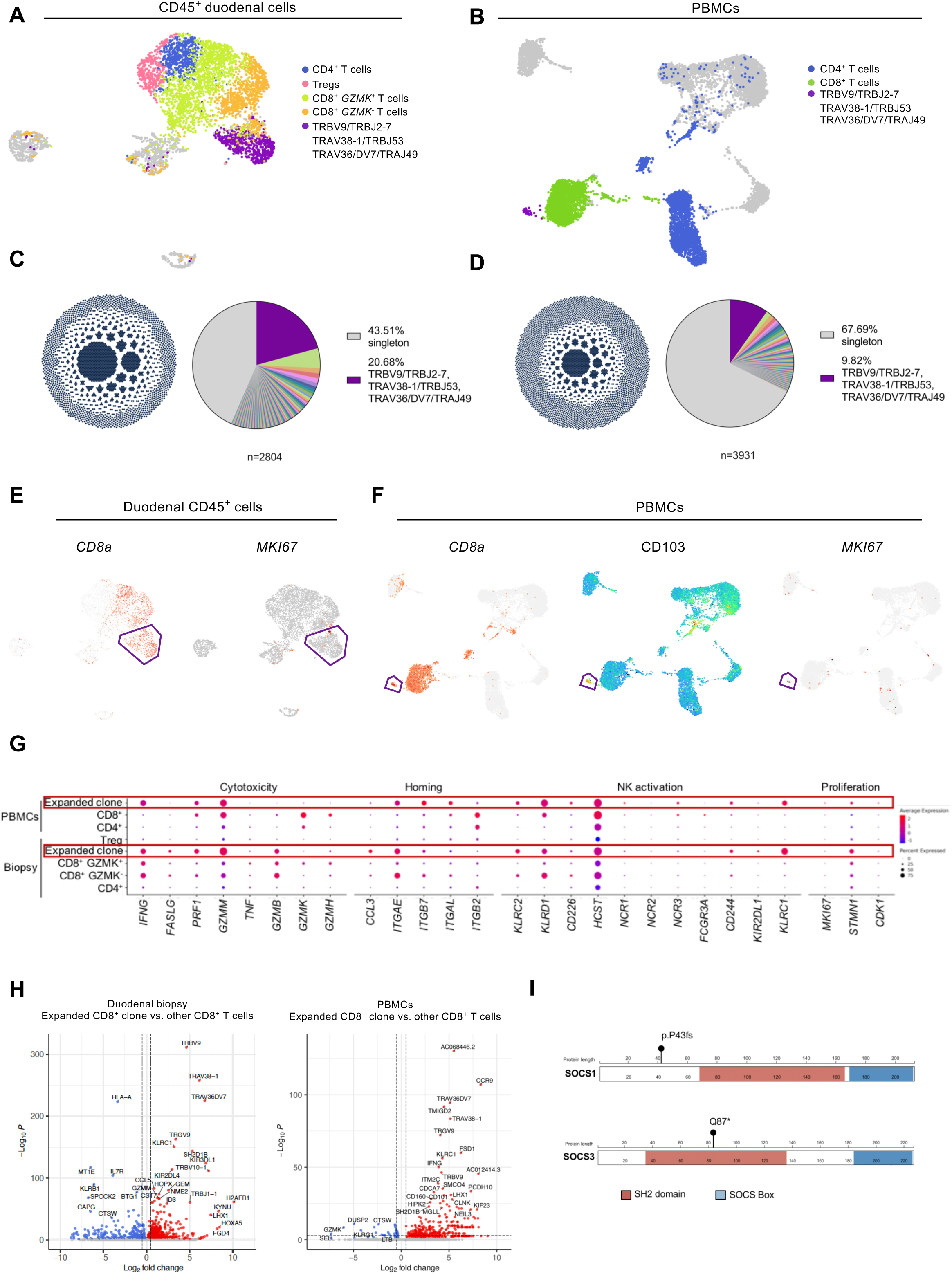
Clonal expansion of CD8⁺ T cells carrying SOCS mutations in RCD1. **A-B.** UMAP of duodenal CD45⁺ cells (A) and PBMCs (B) of patient P10. Dominant CD8⁺ clonotype (TRBV9/TRBJ2-7, TRAV38-1/TRAJ53, TRAV36/DV7/TRAJ49) is highlighted in purple; cells with no TCRαβ expression are shown in grey. **C-D.** Clonal TCRαβ organization in biopsy (n=2,804) and PBMCs (n=3,931). Dots represent single cells, grouped by clonotype. Larger clonotypes appear centrally, smaller ones at the periphery. The dominant clonotype (purple), other expanded clones (color-coded), and singleton cells (grey) are shown. **E-F.** Expression of selected markers: CD8 and MKI67 transcripts, and surface CD103 protein. **G.** Dot plots comparing expression of selected transcripts between indicated T cell subsets. **H.** Volcano plots of genes differentially expressed in the dominant CD8⁺ T cell clone carrying SOCS1 and SOCS3 mutations compared to other CD8⁺ T cells, in the blood (left) and in the duodenal biopsy (right) of RCD1 patient P10 (adjusted p-value<0.05). **I.** Schematic representation of the SOCS1 and SOCS3 mutations identified in the expanded CD8⁺ T-cell clone.

To investigate the genetic basis of this expansion, Vβ1⁺ CD8⁺ T cells and control Vβ1⁻ T cells were sorted from duodenum and blood and analyzed by WES, revealing monoallelic somatic loss-of-function (LOF) mutations in *SOCS1* (c.129delC, p.P43fs) and *SOCS3* (c.259C>T, p.Q87*) (**Figure 6I** and **Table S6**) exclusively in the expanded Vβ1⁺ CD8⁺ T cells. Interestingly, these two genes encode key negative regulators of the JAK-STAT pathway^48^ and inherited monoallelic SOCS1 loss-of-function mutations have been linked to autoimmunity and intestinal inflammation^49,50^.

## Discussion

Here, integrated transcriptomic and genomic analysis of RCD1 and RCD2 provides evidence for a pathophysiological continuum between these conditions. It identifies novel diagnostic and therapeutic targets, while also exposing the challenge of resistance to targeted therapies, particularly JAK inhibitors in RCD2.

RCD2 tumor cells exhibited a distinct transcriptional profile, combining features of ILC1, NK cells, and conventional CD8^+^ IELs. This profile, shared across patients, supports their origin from a rare subset of innate-like sCD3^-^iCD3^+^ IELs found in the normal gut^13^. In addition to NKp46, already recognized as a diagnostic marker, RCD2 tumor cells strongly expressed NK-related transcripts such as SH2D1B and KIR2DL4, and frequently reactivated cancer-testis genes, which may contribute to tumorigenesis or represent potential therapeutic targets^27^. Despite lacking surface TCR/CD3 expression, RCD2 tumor cells harbor clonal TCR rearrangements^12^. In a previous study, we showed that 70% of 28 RCD2 cases exhibited out-of-frame or missing *TCRD*, *G* or *B* rearrangements, preventing surface TCR expression^13^. Here, single-cell full-length V(D)J sequencing confirmed the absence of α or β chain transcripts in most cases. Unexpectedly, two cases displayed multiple atypical α and/or β chain transcripts, suggesting ongoing TCR recombination, a finding recently reported in two other RCD2 cases^17^. As in our study, no RAG1/2 transcripts were detected; however, ongoing TCR recombination has been proposed to promote rapid progression toward aggressive EATL^7,17^. Indeed, aberrant RAG activity is a known driver of chromosomal translocations in lymphoid malignancies, targeting authentic or cryptic recombination signal sequences (RSSs) within immunoglobulin and TCR gene loci ^51–53^.

The presence of *JAK1* and/or *STAT3* GoF mutations in nearly all lymphomas complicating CeD suggests that the JAK/STAT pathway is a possible therapeutic target. However, the highly complex mutational landscape observed in RCD2^15^ may promote ITH and tumor escape. Supporting this concern, transcriptional analysis demonstrated marked ITH increasing with disease progression. Notably, a strong JAK1/STAT3 transcriptional signature was confined to a few cytotoxic subclusters, which may contribute to epithelial damage and be sensitive to JAK1-STAT3 inhibition. In contrast, this signature was absent in the proliferative population, which is present even in low-burdens cases and likely comprises tumor-initiating cells. Taken together, these findings help explain recent clinical observations showing that pan-JAK inhibitors improved histologic lesions in RCD2 patients without reducing tumor cell numbers^19–21^. Further highlighting the risk of selecting RCD2 cells resistant to targeted therapy, *in vitro* treatment of RCD2 cell lines with the JAK inhibitor ruxolitinib led to the emergence of resistant clones carrying mutations that were absent or present at low levels in untreated controls, suggesting selection of pre-existing minor subclones with multiple co-selected variants. Notably, variants with high pathogenicity scores were identified in two tumor suppressor genes, *CDK13* and *RASAL1*^38,39^*. CDK13* encodes a cyclin-dependent kinase, and its inactivating mutations have been shown to promote the oncogenic accumulation and translation of prematurely terminated RNA^38^, while *RASAL1* negatively regulates RAS signaling^39^, a pathway known to contribute to resistance to JAK inibitors^54^. These findings are consistent with drug resistance mechanisms. However, functional validation is needed to confirm the role of these mutations in therapeutic escape.

Unlike RCD2, the mechanisms underlying RCD1 remain unclear. Traditionally considered as a non-clonal, immune-mediated disorder, potentially related to a switch toward gluten-independent autoimmunity^12^, this paradigm is increasingly challenged by several observations. We previously reported association between RCD1 and clonal T-LGL infiltrate of the duodenal mucosa^55^ or indolent intestinal T-cell lymphomas^56^. In this study, one patient initially diagnosed with RCD1 later developed both RCD2 and a CD4^+^ lymphoproliferation, each with a distinct genomic profile, suggesting a distinct clonal origin. In another RCD1 patient, single-cell V(D)J sequencing revealed a large clonal CD8⁺ T cell expansion harboring *de novo* LOF mutations in SOCS1 and SOCS3, two key negative regulators of the JAK-STAT pathway^48^. This clone displayed a cytotoxic and proliferative transcriptomic signature, with NK-like features (e.g;, *SH2D1B*, *KIR2DL4*, *KLRC1*), suggesting enhanced effector function and innate-like reprogramming. Additional signatures, including CD103 surface expression and high *CCR9* transcription in circulating cells, support an intestinal or gut-draining lymph node origin, with subsequent recirculation into the bloodstream. This molecular profile is consistent with a pathologically activated cytotoxic lymphocyte population carrying GOF STAT3 mutations that can drive IL-15 and NKG2D-dependent autoimmune tissue damage ^57^. These findings align with recent observations by Singh et al., who identified similar clonal expansions in 2 of 4 RCD1 patients^17^.

Altogether, these results challenge the binary classification of RCD based on IEL phenotype and clonality. They show that some RCD1 patients can develop clonal lymphocyte expansions with somatic mutations. Similar to those in sCD3⁻iCD3⁺ IELs in RCD2, these mutations may enhance responsiveness to JAK-STAT–activating cytokines and promote clonal persistence in the inflamed mucosa. Clinical differences may reflect distinct cells of origin: sCD3⁺ T IELs in RCD1 versus sCD3⁻iCD3⁺ innate-like IELs in RCD2, as well as differences in mutational burden. Consistent our previous findings^15^, WES revealed multiple oncogenic mutations in RCD2 tumor cells but only few in the expanded RCD1 clone. These data illustrate how chronic inflammation in CeD can drive diverse lymphoproliferative disorders through convergent somatic mutations, particularly those activating JAK1-STAT3, potentially explaining EATL development in CeD or RCD1 patients without prior RCD2.

This study has several limitations. In RCD2, resistance to targeted therapy was demonstrated *ex vivo*, limiting generalizability. In both RCD1 and RCD2, the origin of somatic mutations remains unclear. They may result from clonal hematopoiesis, an age-related process exacerbated by chronic inflammation and linked to lymphoid malignancies including T-LGL leukemia^58^, or arise during inflammation-driven lymphocyte proliferation, as observed in autoimmune diseases like rheumatoid arthritis and Sjögren’s syndrome, where B-cell mutations accumulate and promote lymphomagenesis^59^. Finally, while RCD2 IELs are known to exert NK-like cytotoxicity against enterocytes^36,60,61^, the role of clonal T lymphocytes in epithelial damage in RCD1 remains unclear. Whether these cells are autoreactive, similar to pathogenic B cells in rheumatoid arthritis, is an open question with potential therapeutic relevance.

In conclusion, our findings provide new insights into mechanisms of GFD refractoriness in CeD and highlight the need for molecularly guided therapies. More broadly, they suggest that clonal T-cell expansions with cytokine-sensitizing mutations may contribute to treatment resistance in other chronic inflammatory diseases, such as Crohn’s disease, where post-operative relapses have been linked to oligoclonal T-cell expansions^62^.

## Supporting information

Supplementary references

## Acknowledgments

The authors are supported by institutional grants from INSERM and Université Paris Cité and by grants from Plan Cancer Aviesan-SCILD 2019-2022, INCa (PLBIO-2022-098), Agence Nationale pour la Recherche (ANR-24-CE17-5708), Foundation Princesse Grace de Monaco, ERC Marie Skłodowska-Curie Individual Fellowships SingCelCD, 843042) and Association Française Des Intolérants au Gluten (AFDIAG). Institute Imagine is supported by the Investissement d’Avenir grant ANR-10-IAHU-01. We thank Dr. Lilia Ladada for her valuable assistance in the analysis and interpretation of exome variants identified in ruxolitinib-resistant cell lines using the MobiDetails platform.

