## Supplemental Figures for "CLONAL LYMPHOCYTE EXPANSIONS AND JAK-STAT PATHWAY MUTATIONS DEFINE A PATHOGENIC CONTINUUM DRIVING RESISTANCE TO GLUTEN-FREE DIET IN CELIAC DISEASE"

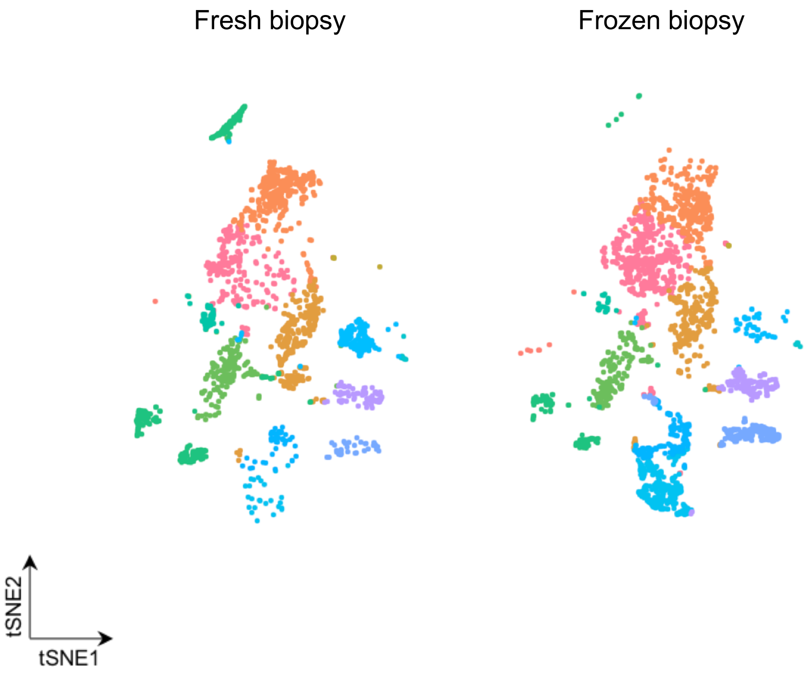


**Supplementary Figure 1**: **Comparison of the transcriptional profiles between fresh and frozen biopsy samples.** tSNE plot illustrating clustering of 2,548 cells from frozen duodenal biopsies, 1,935 cells from fresh biopsies, and 3,135 PBMCs. Minimal differences observed between transcriptional profiles of fresh and frozen samples validate cryopreservation methods.

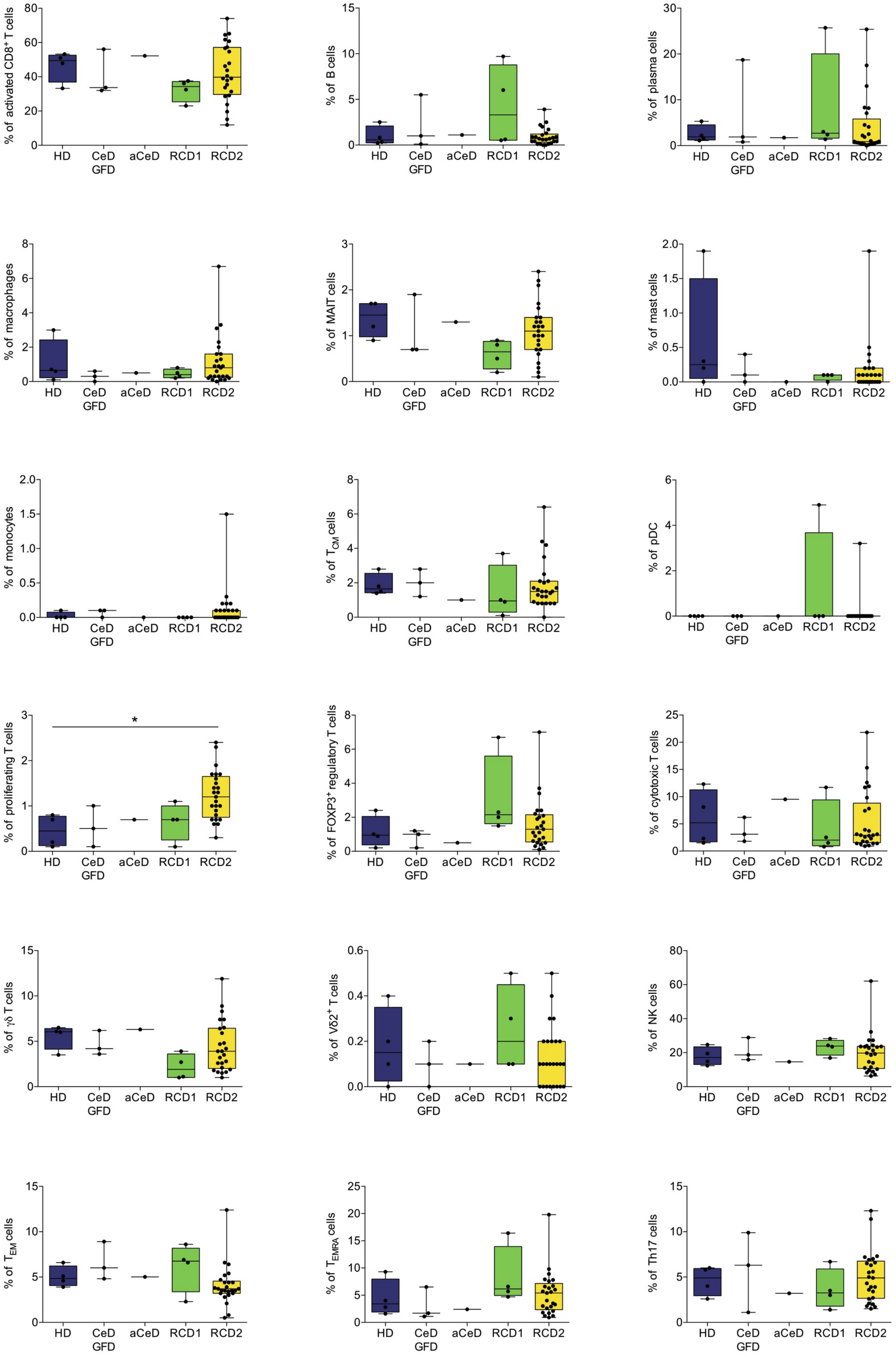


**Supplementary Figure 2: Immune subtypes across RCD2 and control samples.** Bar plot showing the percentages of the different immune cell types identified by sc-RNAseq in RCD2, RCD1, aCeD, GFD, and controls.

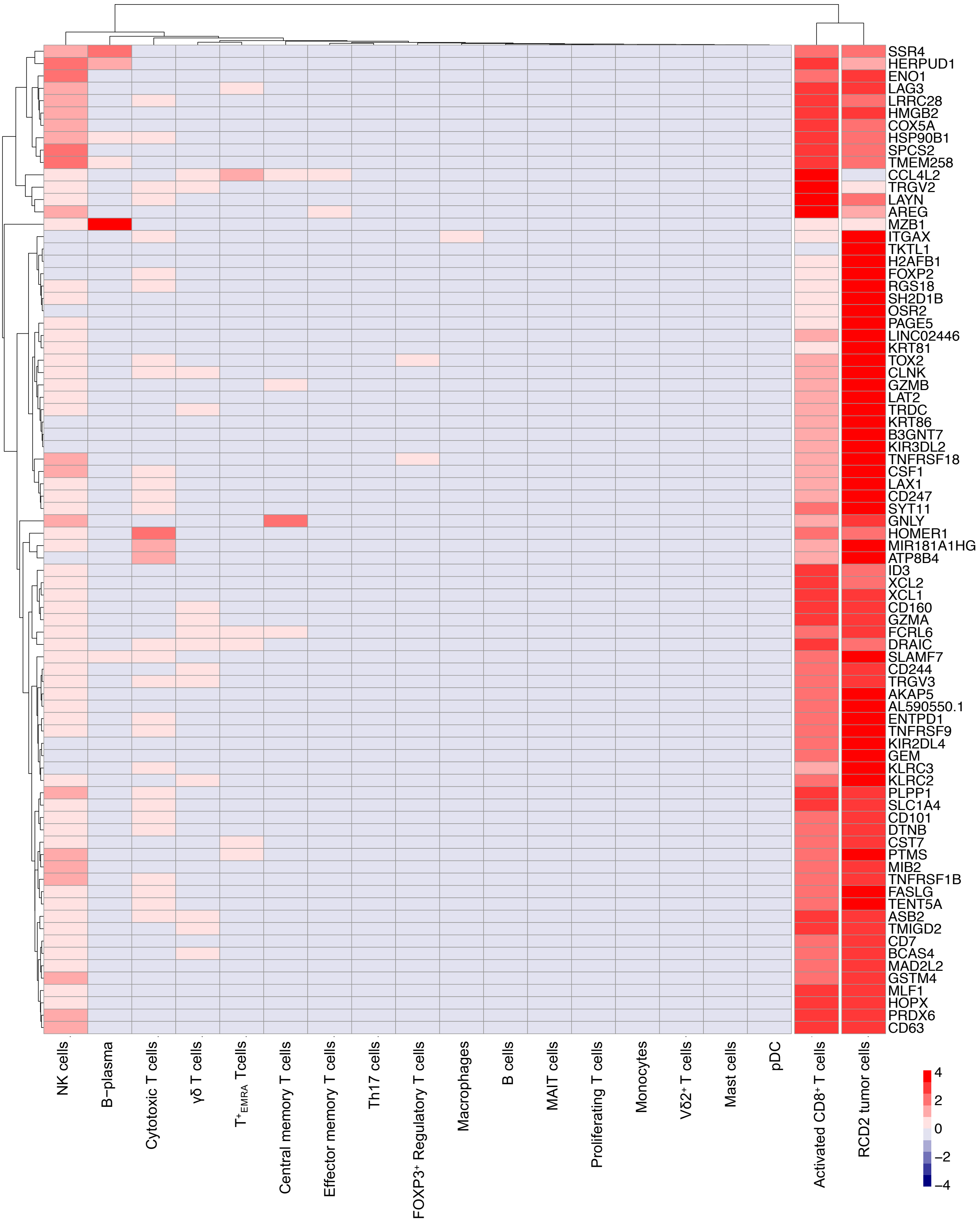


**Supplementary Figure 3. Comparison of the transcriptomic profile of RCD2 cells and other immune cell types in duodenal biopsies**

Heatmap illustrating the expression of overexpressed transcripts in RCD2 cells across various immune cell types identified in duodenal biopsies by scRNAseq.

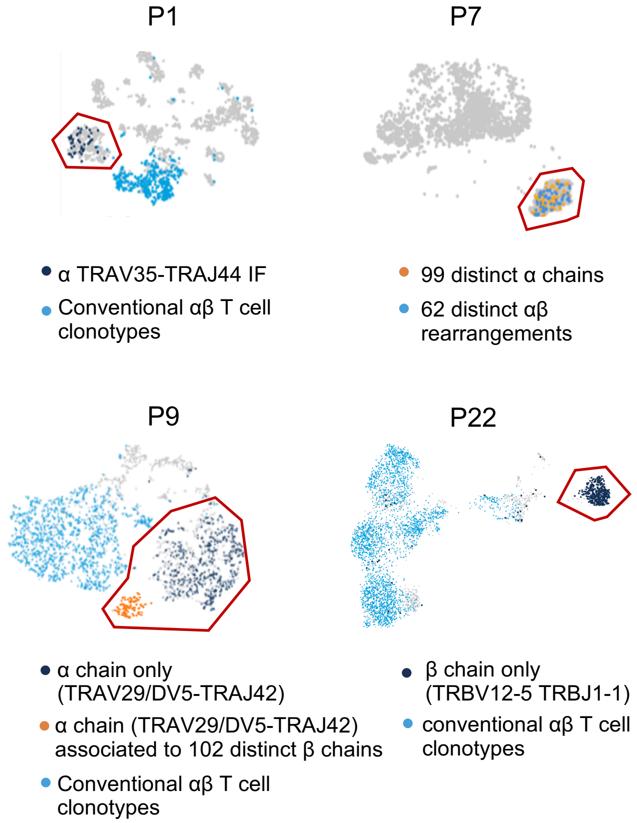


**Supplementary Figure 4: Expression of atypical TCR α and β chain transcripts in RCD2 patients**
In 6 out of 13 RCD2 cases, single cell V(D)J immunoprofiling of cells isolated from frozen intestinal biopsies reveals α and/or β chain transcripts in malignant RCD2 cells (circled in red). In 2 cases, RCD2 cells harbored a unique α chain but no β chain (ex P1). In one case (P7), out of 2777 tumor cells expressing a $V\gamma5^{+}$ transcript, 99 displayed an α chain transcript (57 distinct) and 62 showed both α and β chains transcripts, all of which were distinct. In one case (P9), RCD2 cells harbored a unique α chain associated or not with multiple β chains. In 2 other cases tumor cells harbored a unique β chain but no α chain (ex P22). No transcript encoding RAG was detected.

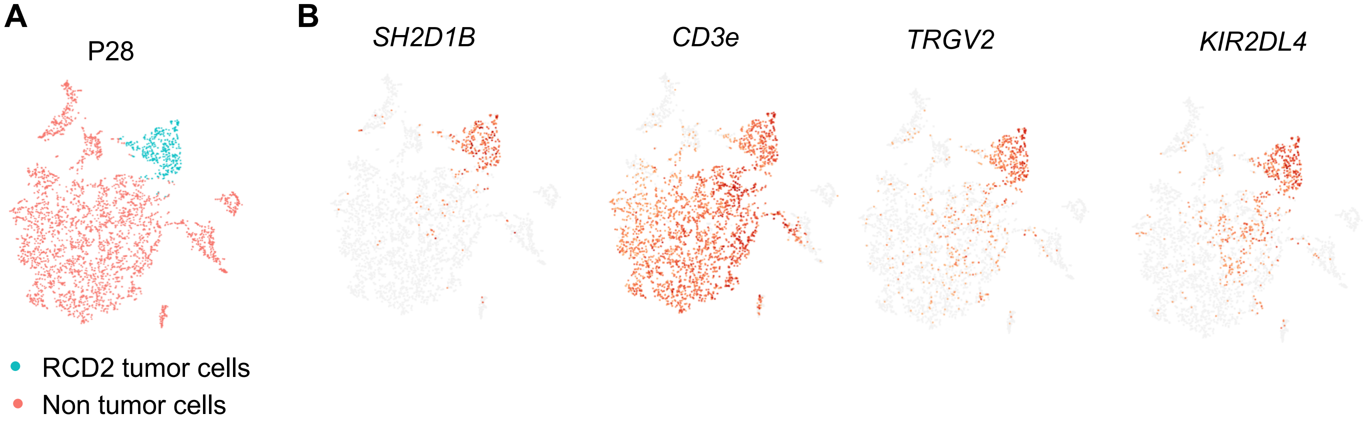


**Supplementary Figure 5: Transcriptomic profile of TCRγδ⁺ tumor cells in RCD2 patient P28**
**A.** tSNE plot illustrating tumor cells displaying surface CD3 and TCRγδ in one RCD2 patient (P28). **B.** The transcriptomic profile was largely consistent with other RCD2 samples, showing high expression of *SH2D1B* and *KIR2DL4* along with uniformly strong expression of *TRGV2* transcripts.

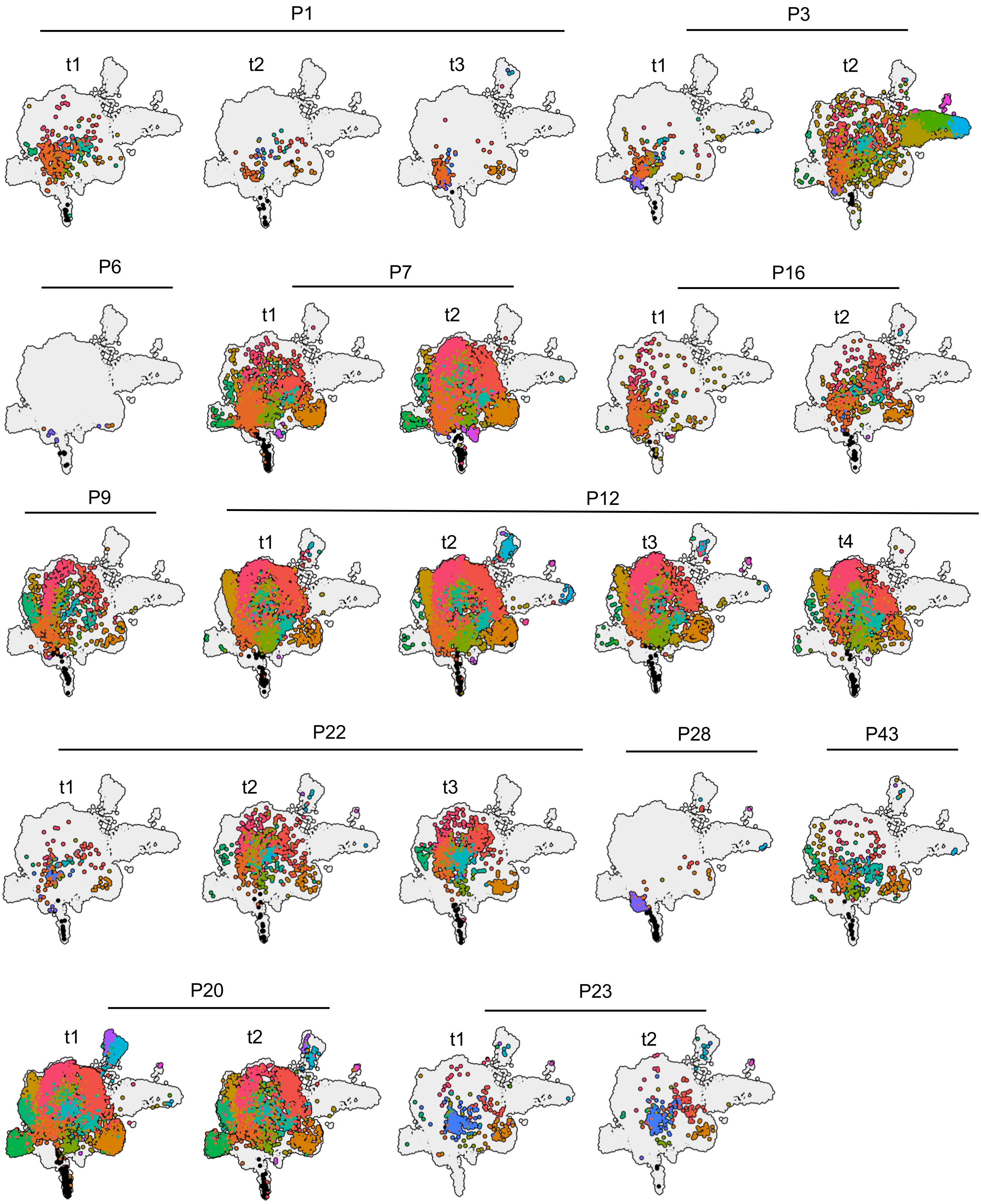


**Supplementary Figure 6: UMAP visualization of RCD2 tumor cells across 24 individual duodenal biopsies**
UMAP plots showing the distribution of RCD2 tumor cell clusters across individual duodenal biopsies, organized by patient and time point. Cells from different clusters are projected onto each map to illustrate temporal and inter-patient heterogeneity.

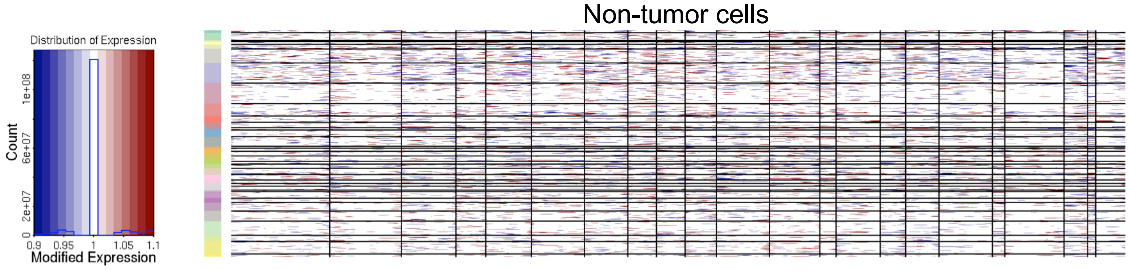


**Supplemental Figure 7: CNV analysis in non-tumor control cells from duodenal biopsies**

Inferred CNV profiles from all non-tumor duodenal $CD45^+$ cells (**see Figure S7**), including those from the same samples as well as from additional control biopsies (n = 5 controls; n = 5 CeD patients on a gluten-free diet). The y-axis represents individual cells, and the x-axis corresponds to genomic position.

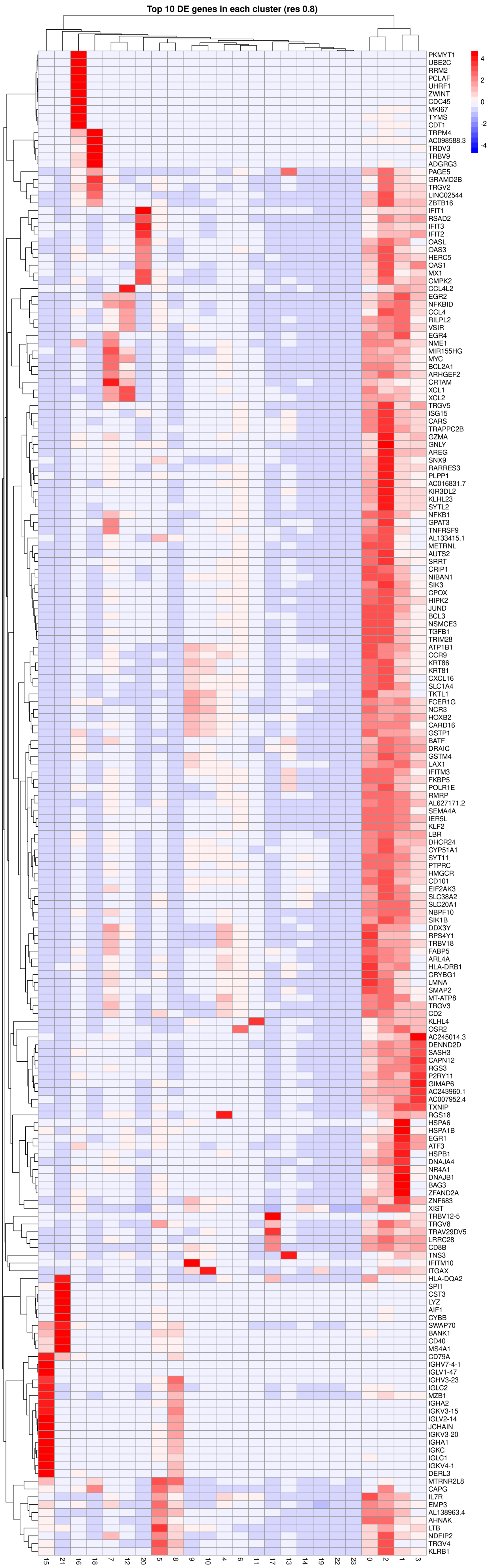


**Figure S8 : Top 10 differentially expressed genes across RCD2 tumor cell clusters**

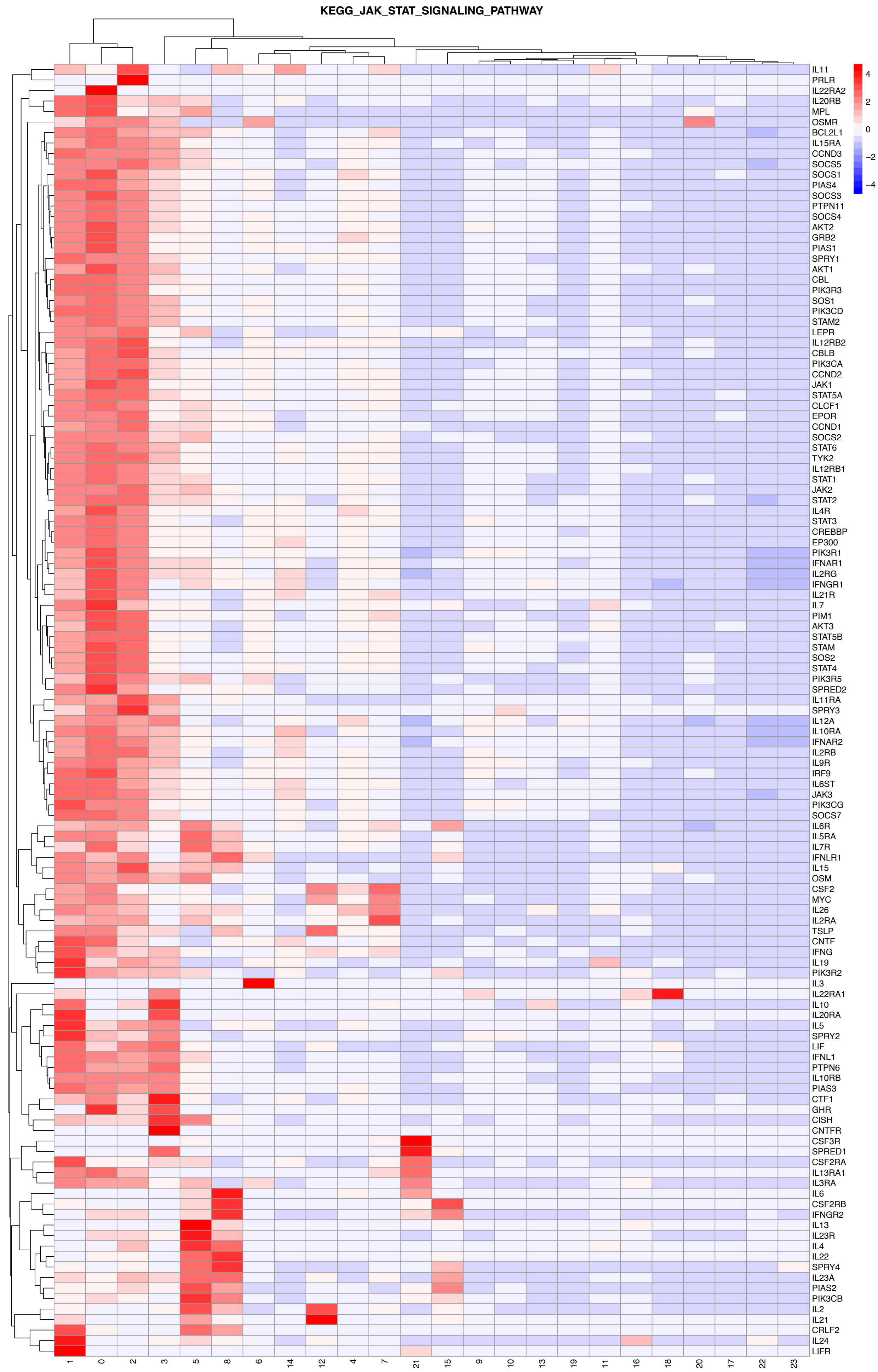


**Figure S9 : Heterogeneous expression of JAK-STAT pathway genes across RCD2 tumor cell clusters**

**Supplementary Figure 9: Heterogeneous expression of JAK-STAT pathway genes across RCD2 tumor cell clusters.** Heatmap showing the scaled expression of genes from the KEGG JAK-STAT signaling pathway across tumor cell clusters identified by unsupervised analysis in RCD2 biopsies. Each column represents a tumor cell cluster, and each row corresponds to a pathway gene. Expression levels are Z-score normalized, with red indicating higher expression relative to the mean and blue indicating lower expression. The observed heterogeneity highlights variable engagement of the JAK-STAT pathway among tumor subpopulations, with clusters 0, 1, and 2 showing elevated expression of components such as *JAK1*, *STAT3*, and *SOCS1*, and cluster 16 exhibiting notably lower activation.

**A**

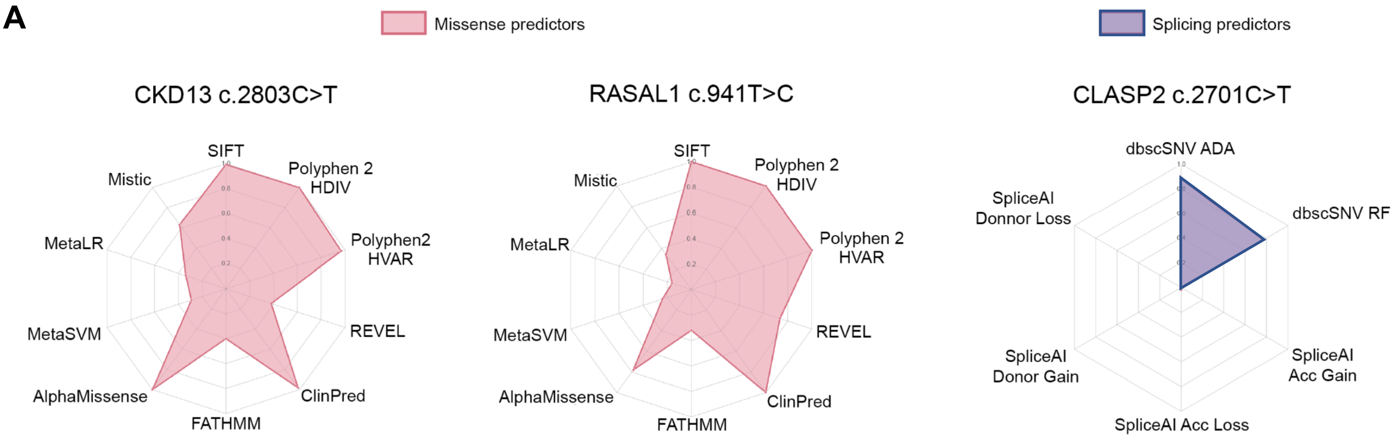


**B**

| | ACMG classification for gene mutated in 5/6 cell line minimum | | | | | | | | |
|---|---|---|---|---|---|---|---|---|---|
| | CLCA2 | CDK13 | RASAL1 | DENND2B | RNF149 | CLASP2 | STK11IP | ADGRB1 | HIVEP3 |
| hg38 InterVar | Uncertain | Uncertain | Uncertain | Uncertain | | Pathogenic | Uncertain | Uncertain | Uncertain |
| | | PM1 | PM1 | PM1 | | PVS1 | | | PM1 |
| | PM2 | PM2 | PM2 | PM2 | | PM2 | PM2 | PM2 | PM2 |
| | PP3 | PP2 | | | | PP3 | | PP3 | |
| GeneBe | Uncertain | Uncertain | Uncertain | Uncertain | Likely begnign | Uncertain | Uncertain | Uncertain | Uncertain |
| | | PM1 | | | | | | | |
| | PM2 | PM2 | PM2 | PM2 | PM2 | PM2 | PM2 | PM2 | PM2 |
| | PP3_Moderate | PP3 | PP3_Moderate | | | | | | |
| | | | | BP4 | BP4_Strong | | | | |

| | ZNF829 | PRKCQ | ZNF214 | ZNF140 | AGBL5 | DNAH11 | TRIM48 | SYNE2 | LARP6 |
|---|---|---|---|---|---|---|---|---|---|
| hg38 InterVar | Uncertain | | Uncertain | Uncertain | Uncertain | | Uncertain | Uncertain | |
| | PM1 | | PM1 | PM1 | PM1 | | | PM1 | |
| | PM2 | | PM2 | PM2 | PM2 | | PM2 | | |
| | | | | | PP2 | | | | |
| | | | | | BP1 | | | | |
| | | | BP4 | BP4 | | | BP4 | BP4 | |
| GeneBe | Uncertain | | Uncertain | Uncertain | Uncertain | Likely begnign | Uncertain | Benign | |
| | | | | | | PM2 | | | |
| | | | | | | BP4_Strong | | BP4_Strong | |
| | | | | | | | | BS2 | |

| | |
|---|---|
| PVS1 | Null variant (nonsense, frameshift, canonical +-2 splice sites, initiation codon, single or multiexon deletion) in a gene where LOF is a known mechanism of disease |
| PM1 | Located in a mutational hot spot and/or critical ans well-established functional domain (e.g., active site of an enzyme) without benign variation |
| PM2 | Absent from controls (or at extremely low frequency if recessive) in Exome Sequencing Project, 1000 Genomes Project, or Exome Aggregation Consortium |
| PP2 | Missense variant in a gene that has a low rate of benign missense variation and in which missense variants are a common mechanism of disease |
| PP3 | Multiple lines of computational evidence support a deleterious effect on the gene or gene product (conservation, evolutionary, splicing impact, etc.) |
| PP3 Moderate | Multiple lines of computational evidence support a deleterious effect on the gene or gene product (conservation, evolutionary, splicing impact, etc.) |
| BP1 | Missense variant in a gene for which primarily truncating variants are known to cause disease |
| BP4 | Multiple lines of computational evidence suggest no impact on gene or gene product (conservation, evolutionary, splicing impact, etc.) |
| BP4 Strong | Multiple lines of computational evidence suggest no impact on gene or gene product (conservation, evolutionary, splicing impact, etc.) |
| BS2 | Observed in a healthy adult individual for a recessive (homozygous), dominant (heterozygous) disorder, with full penetrance expected at an early age |

**Supplementary Figure 10 : *In silico* pathogenicity prediction of candidate resistance variants.**

**Supplementary Figure 10:** ***In silico*** **pathogenicity prediction of candidate resistance variants.**
Variants were analysed in mobidetails plateform[89] when possible. **A.** Left panel: Radar chart displaying missense variant predictors for CDK13 c.2803C>T and RASAL1 c.941T>C variants. Values are normalised (0-1), where 0 indicates the least damaging prediction and 1 the most damaging for each predictor. Right panel: Radar chart showing splicing effects predictors for the CLASP2 c.2701C>T variant. Values are normalised as above. **B.** Semi-automated classification of variant pathogenicity according to the American College of Medical Genetics and Genomics (ACMG) guidelines was performed using the interVar tool[90] (reported as "hg38 InterVar" in the table), and the GeneBe portal[91], which compiles data from multiple sources to support ACMG-based annotation[91] (reported as "GeneBE" in the table).

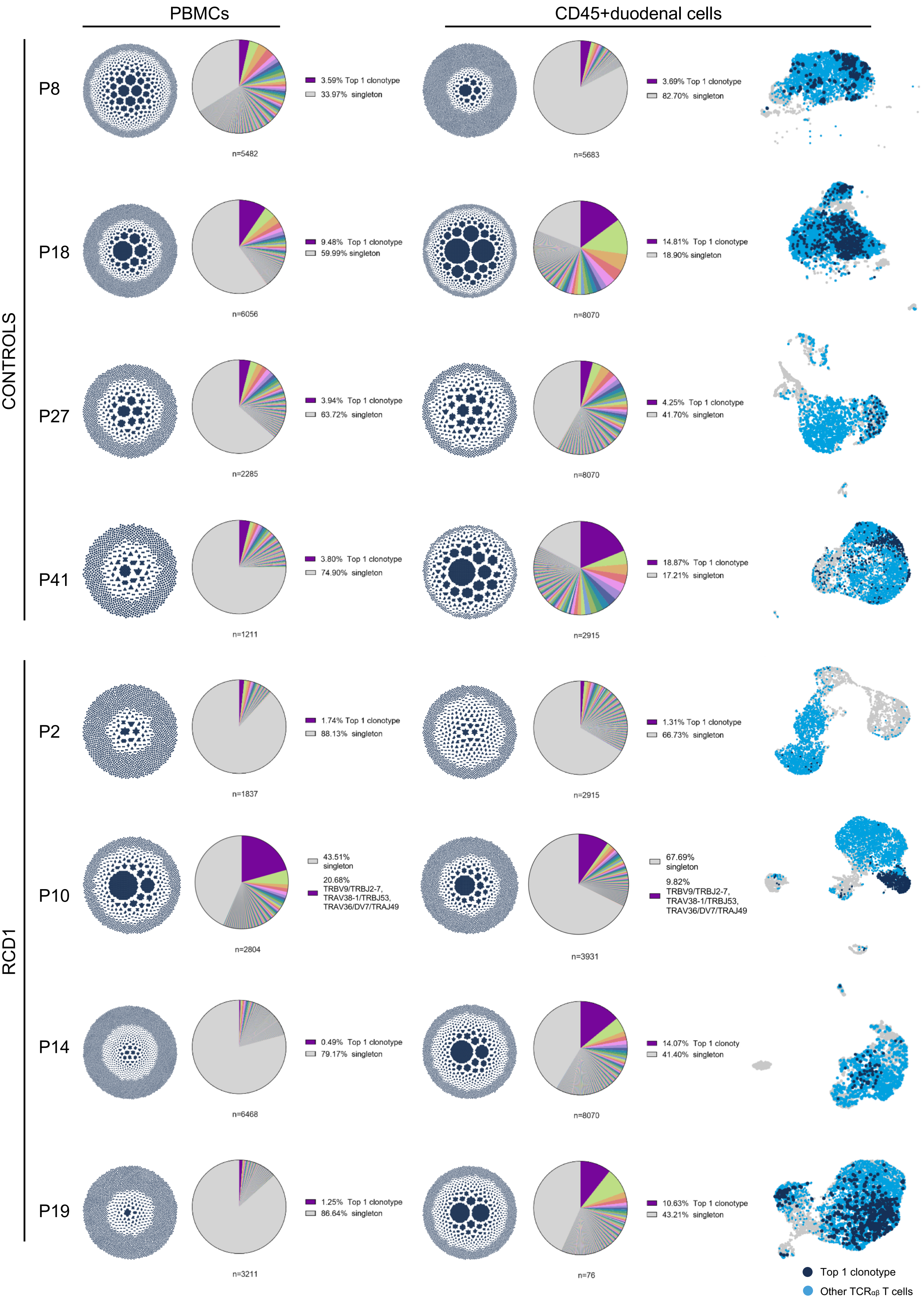


**Supplementary Figure 11: Clonal organization and distribution of paired TCRαβ sequences in RCD1 and control patients' biopsies and PBMCs**

**Supplementary Figure 11: Clonal organization and distribution of paired TCRαβ sequences in RCD1 and control patients' biopsies and PBMCs**

Clonal organization of paired TCR α and β sequences generated by the Loupe VDJ Browser. The distribution plot displays cells grouped by clonotypes, with each dot representing a single cell (10x barcode). Barcodes are grouped by clonotype ID, and white spaces separating different clonotypes. Larger clonotypes are positioned centrally, while smaller clonotypes are at the periphery. The proportion of cells in the indicated cell types with assigned paired TCRα and TCRβ sequences is shown for biopsies and PBMCs from 4 controls and 4 RCD1. The most expanded clonotype (purple) is highlighted, along with other expanded clones (color-coded) and unique singletons (grey). UMAP of duodenal $CD45^+$ cells from 4 controls and 4 RCD1. Cells belonging to the top one clonotype are highlighted in dark blue while other TCRαβ T cells are in light blue. Unexpectedly, controls P18 and P41, despite having normal histology, exhibited an expanded clonotype—possibly related to unexplained chronic diarrhea and a family history of autoimmune disease in patient P41.

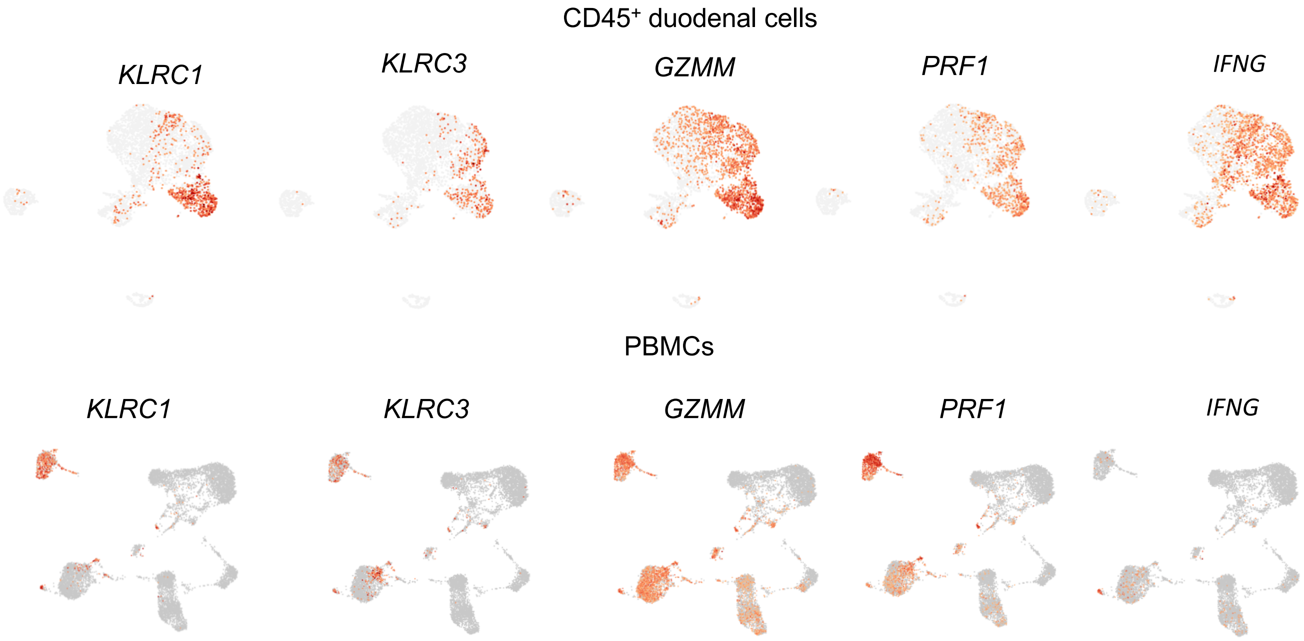


**Supplementary Figure 12: Activation and cytotoxic gene expression in CD45$^{+}$ duodenal cells and PBMCs of RCD1 P10.**

UMAP showing normalized expression levels of *KLRC1*, *KLRC3*, *GZMM*, *PRF1* and *IFNG* transcripts in duodenal CD45$^{+}$ cells and PBMCs from RCD1 patient P10.

| | Clinical Status at Sampling | | | | | | Diagnostic Features | | | | | | |
|---|---|---|---|---|---|---|---|---|---|---|---|---|---|
| | | | | | | | Histology/IHC | | | | Flow cytometry | | |
| | patient ID | time point | age | gender | Strict GFD since CeD diagnosis | Treatment | EATL | villous atrophy (duodenum) | Marsh score | IEL/100EC | % RCD2 IEL/CD45$^+$ | RCD2 LPL/CD45$^+$ | RCD2 PBL/CD45$^+$ |
| RCD1 (n=4) | P2 | t1 | 34 | F | yes | no | no | yes | IIIa | 50 | NA | NA | NA |
| | P19 | t1 | 59 | F | yes | budesonide 6mg/day | no | yes | IIIB | 40 | NA | NA | NA |
| | P10 | t1 | 72 | F | yes | no | no | yes | IIIB | 1 | NA | NA | NA |
| | P14 | t1 | 64 | M | yes | no | no | yes | IIIa | 40 | NA | NA | NA |
| RCD2 (n=24) | P1 | t1 | 43 | F | yes | budesonide 3mg/day | no | no | II | 80 | 66% | 7% | <1% |
| | | t2 | 44 | F | yes | budesonide 6mg/day; MTX 15mg/week | no | yes | IIIa | 80 | 66% | 25% | 0.05% |
| | | t3 | 46 | F | yes | budesonide 9mg/day | yes | yes | IIIa | 50 | 80% | 28% | <0,03% |
| | P3 | t1 | 78 | F | yes | budesonide 9mg/day | no | yes | II | 40 | ND | ND | 0,10% |
| | | t2 | 79 | F | yes | budesonide 6mg/day | yes | no | II | 50 | ND | ND | ND |
| | P6 | t1 | 69 | M | yes | budesonide 9mg/day | no | yes | IIIA | 50 | 67% | 18% | 4% |
| | P7 | t1 | 39 | F | yes | budesonide 9mg/day | no | no | II | 70 | 83% | 21% | 1% |
| | | t2 | 42 | F | yes | no | no | yes | IIIA | 50 | 80% | 68% | <1% |
| | P9 | t1 | 75 | F | yes | budesonide 9mg/day | yes | yes | IIIB | 80 | 77% | 4% | 0% |
| | P12 | t1 | 62 | M | no | budesonide 9mg/day | yes | yes | IIIA | ND | ND | ND | ND |
| | | t2 | 63 | M | no | budesonide 9mg/day | yes | yes | IIIA | ND | 99% | 58% | 1% |
| | | t3 | 63 | M | no | budesonide 9mg/day | yes | yes (Ileum) | IIIA | >30 | ND | ND | ND |
| | | t4 | 64 | M | no | prednisone 5mg/day | yes | yes | IIIC | 70 | 95% | 58% | 0,50% |
| | P16 | t1 | 65 | M | yes | prednisone 5mg/day | no | no | II | 40 | 25% | 9% | <1% |
| | | t2 | 66 | M | yes | prednisone 5mg/day | no | no | II | 30 | 22% | 10% | 0,03% |
| | P20 | t1 | 74 | F | yes | no | no | no | II | 70 | 93% | 36% | 3% |
| | | t2 | 76 | F | yes | budesonide 9mg/day | no | no | IIIA | 40 | 96% | 43% | 0,02% |
| | P22 | t1 | 55 | M | no | budesonide 9mg/day | yes | yes | IIIA | 20 | 36% | 11% | 1% |
| | | t2 | 55 | M | no | budesonide 9mg/day | yes | yes | IIIA | ND | 80% | 15% | 0,10% |
| | | t3 | 56 | M | no | budesonide 9mg/day | yes | yes | IIIA | >50 | 73% | 17% | 0,00% |
| | P23 | t1 | 45 | F | yes | budesonide 9mg/day | no | yes | IIIA | 100 | 51% | 15% | 0,10% |
| | | t2 | 46 | F | yes | no | no | yes | IIIA | 70 | 38% | 8% | 0,04% |
| | P28 | t1 | 56 | F | yes | no | no | yes | IIIA | 70 | 81% | 11% | 0.4% |
| | P43 | t1 | 73 | F | yes | no | no | yes | IIIA | 40 | 37% | 11% | 3% |
| HD (n=4) | P8 | t1 | 48 | M | no | no | no | no | I | ND | NA | NA | NA |
| | P18 | t1 | 48 | M | no | no | no | no | I | ND | NA | NA | NA |
| | p27 | t1 | 82 | F | no | no | no | no | I | ND | NA | NA | NA |
| | P33 | t1 | 56 | F | no | no | no | no | I | ND | NA | NA | NA |
| CeD GFD (n=5) | P24 | t1 | 29 | F | yes | no | no | no | I | ND | NA | NA | NA |
| | P25 | t1 | 39 | F | yes | no | no | yes | IIIA | 70 | NA | NA | NA |
| | P37 | t1 | 39 | F | yes | budesonide 3mg/day | no | no | I | ND | NA | NA | NA |
| | P40 | t1 | 52 | F | yes | no | no | no | I | ND | NA | NA | NA |
| | P47 | t1 | 51 | F | yes | no | no | no | I | 20 | NA | NA | NA |
| aCeD (n=1) | P42 | t1 | 38 | M | no | no | no | yes | IIIA | ND | NA | NA | NA |

**Supplementary Table S1. Clinical, histologic, and immunologic features of patients with CeD, RCD1, and RCD2.** In flow cytometry analysis, RCD2 cells are defined as $iCD3\varepsilon^+CD103^+$ cells lacking surface expression of $CD3\varepsilon$. In patient P28, RCD2 cells correspond to $CD3^+TCR\gamma\delta^+$ cells.

| Type | ID1 | chr | Amplicon_start | Amplicon_end | Mutation | Gene |
|---|---|---|---|---|---|---|
| Region | chr7:48547631-48547631 | chr7 | 48547395 | 48547658 | p.F4504V | ABCA13 |
| Region | chr7:152513614-152513614 | chr7 | 152513509 | 152513779 | p.R161C | ACTR3B |
| Region | chr4:74306339-74306339 | chr4 | 74306105 | 74306375 | p.E97D | AFP |
| dbSNP | chr6:138197104-138197303 | chr6 | 138197080 | 138197330 | CNV | TNFAIP3 |
| dbSNP | chr6:138198182-138198384 | chr6 | 138198155 | 138198405 | CNV | TNFAIP3 |
| Region | chr12:96354369-96354369 | chr12 | 96354146 | 96354388 | p.G261R | AMDHD1 |
| Region | chr4:114120223-114120223 | chr4 | 114120013 | 114120246 | p.K93N | ANK2 |
| Region | chr10:61819189-61819189 | chr10 | 61818949 | 61819219 | SpliceAccDon | ANK3 |
| Region | chr11:120317130-120317130 | chr11 | 120316962 | 120317152 | p.L436P | ARHGEF12 |
| Region | chr12:123465040-123465040 | chr12 | 123464820 | 123465077 | p.A36T | ARL6IP4 |
| Region | chr6:101127592-101127592 | chr6 | 101127372 | 101127614 | p.R730G | ASCC3 |
| Region | chr20:31021233-31021233 | chr20 | 31020988 | 31021258 | p.R411H | ASXL1 |
| Region | chr13:26411314-26411314 | chr13 | 26411089 | 26411335 | p.L923S | ATP8A2 |
| Region | chr3:165548298-165548298 | chr3 | 165548060 | 165548330 | p.R175M | BCHE |
| Region | chrX:39934292-39934292 | chrX | 39934042 | 39934312 | p.E103* | BCOR |
| Region | chr1:49227121-49227121 | chr1 | 49226883 | 49227150 | p.N83T | BEND5 |
| Region | chr10:60558225-60558225 | chr10 | 60557979 | 60558249 | p.L478R | BICC1 |
| Region | chrX:154300601-154300601 | chrX | 154300357 | 154300627 | SpliceAccDon | BRCC3 |
| Region | chr9:121930350-121930350 | chr9 | 121930102 | 121930372 | p.N433T | BRINP1 |
| Region | chr1:190067306-190067306 | chr1 | 190067061 | 190067331 | p.P715S | BRINP3 |
| Region | chr17:38320125-38320125 | chr17 | 38319877 | 38320147 | p.P393S | CASC3 |
| Region | chr7:116140441-116140441 | chr7 | 116140191 | 116140461 | p.A93G | CAV2 |
| Region | chr3:105439096-105439096 | chr3 | 105438853 | 105439123 | SpliceAccDon | CBLB |
| Region | chr6:99997452-99997452 | chr6 | 99997229 | 99997499 | SpliceAccDon | CCNC |
| Region | chr5:24509747-24509747 | chr5 | 24509640 | 24509830 | p.E395G | CDH10 |
| Region | chr5:176013842-176013842 | chr5 | 176013594 | 176013862 | p.R952P | CDHR2 |
| Region | chr3:74413657-74413658 | chr3 | 74413440 | 74413685 | p.K391X | CNTN3 |
| Region | chr1:65656300-65656512 | chr1 | 65656279 | 65656530 | CNV | CNV |
| dbSNP | rs4325172 | chr1 | 65857901 | 65858171 | CNV | CNV |
| dbSNP | rs71244118 | chr1 | 143743262 | 143743532 | CNV | CNV |
| dbSNP | rs111903322 | chr1 | 150040520 | 150040789 | CNV | CNV |
| dbSNP | rs2228099 | chr1 | 150808639 | 150808909 | CNV | CNV |
| dbSNP | rs113314506 | chr1 | 173556687 | 173556932 | CNV | CNV |
| dbSNP | rs7515820 | chr1 | 205553846 | 205554116 | CNV | CNV |
| dbSNP | rs113478407 | chr1 | 212911569 | 212911820 | CNV | CNV |
| dbSNP | rs12731746 | chr1 | 235392301 | 235392571 | CNV | CNV |
| dbSNP | rs2362587 | chr1 | 246021715 | 246021967 | CNV | CNV |
| dbSNP | rs3825175 | chr12 | 122079191 | 122079461 | CNV | CNV |
| dbSNP | rs28651018 | chr12 | 122186109 | 122186299 | CNV | CNV |
| dbSNP | rs3741593 | chr12 | 122261273 | 122261518 | CNV | CNV |
| dbSNP | rs28629903 | chr12 | 122496775 | 122497023 | CNV | CNV |
| dbSNP | rs113128227 | chr12 | 122690789 | 122691042 | CNV | CNV |
| dbSNP | rs3733558 | chr4 | 121957843 | 121958113 | CNV | CNV |
| dbSNP | rs111326362 | chr4 | 123328872 | 123329103 | CNV | CNV |
| dbSNP | rs113388762 | chr4 | 108614889 | 108615159 | CNV | CNV |
| dbSNP | rs111366480 | chr4 | 106158127 | 106158397 | CNV | CNV |
| dbSNP | rs112767825 | chr4 | 107153935 | 107154188 | CNV | CNV |
| dbSNP | rs112234514 | chr4 | 120439991 | 120440261 | CNV | CNV |
| dbSNP | rs113637606 | chr4 | 122250345 | 122250600 | CNV | CNV |
| dbSNP | rs112838003 | chr4 | 122773948 | 122774218 | CNV | CNV |
| sbSNP | rs113888128 | chr4 | 119951511 | 119951781 | CNV | CNV |
| Region | chr1:65306806-65307022 | chr1 | 65306792 | 65307042 | CNV | JAK1 |
| Region | chr1:65310389-65310574 | chr1 | 65310368 | 65310599 | CNV | JAK1 |
| Region | chr1:65311137-65311327 | chr1 | 65311116 | 65311351 | CNV | JAK1 |
| Region | chr1:65312196-65312419 | chr1 | 65312178 | 65312439 | CNV | JAK1 |
| Region | chr1:65325742-65325945 | chr1 | 65325719 | 65325969 | CNV | JAK1 |
| Region | chr1:65330457-65330655 | chr1 | 65330432 | 65330682 | CNV | JAK1 |
| Region | chr1:65339005-65339206 | chr1 | 65338983 | 65339229 | CNV | JAK1 |
| Region | chrX:107431248-107431248 | chrX | 107431017 | 107431275 | p.G533W | COL4A6 |
| dbSNP | rs112949141 | chr6 | 135510777 | 135511047 | CNV | CNV |
| dbSNP | rs62431285 | chr6 | 136590376 | 136590644 | CNV | CNV |
| Region | chr4:1222061-1222061 | chr4 | 1221844 | 1222091 | p.F89Y | CTBP1 |
| dbSNP | rs9321567 | chr6 | 137026197 | 137026465 | CNV | CNV |
| dbSNP | rs648396 | chr6 | 138413052 | 138413290 | CNV | CNV |
| Region | chr10:104591282-104591282 | chr10 | 104591033 | 104591303 | p.P409Q | CYP17A1 |
| dbSNP | rs113062048 | chr6 | 138724576 | 138724846 | CNV | CNV |
| Region | chrX:41203345-41203345 | chrX | 41203104 | 41203369 | p.R276S | DDX3X |
| dbSNP | rs113455315 | chr6 | 139100770 | 139101032 | CNV | CNV |
| Region | chr12:51072560-51072560 | chr12 | 51072333 | 51072598 | p.L339F | DIP2B |
| Region | chr16:20966181-20966181 | chr16 | 20965970 | 20966206 | p.C3675W | DNAH3 |
| Region | chr3:137787123-137787123 | chr3 | 137786937 | 137787167 | p.E568K | DZIP1L |
| Region | chr11:62295105-62295105 | chr11 | 62294883 | 62295153 | p.E2262* | AHNAK |
| Region | chr13:78477351-78477351 | chr13 | 78477116 | 78477386 | p.Y337* | EDNRB |
| Region | chr8:22548678-22548678 | chr8 | 22548432 | 22548702 | p.V120M | EGR3 |
| Region | chr9:75545794-75545794 | chr9 | 75545555 | 75545815 | SpliceAccDon | ALDH1A1 |
| Region | chr22:37770445-37770445 | chr22 | 37770228 | 37770467 | p.F377Y | ELFN2 |
| Region | chr1:79385902-79385902 | chr1 | 79385670 | 79385928 | p.V476D | ELTD1 |
| Region | chr12:132530360-132530360 | chr12 | 132530113 | 132530383 | p.V2372G | EP400 |
| Region | chr8:95686611-95686611 | chr8 | 95686371 | 95686641 | | ESRP1 |
| Region | chr5:150925263-150925263 | chr5 | 150925018 | 150925288 | p.A1809S | FAT2 |
| Region | chr1:152286227-152286227 | chr1 | 152285995 | 152286248 | p.S379R | FLG |
| Region | chr9:132757205-132757205 | chr9 | 132757045 | 132757241 | p.Q19H | FNBP1 |
| Region | chr2:186668776-186668776 | chr2 | 186668581 | 186668800 | p.D5004H | FSIP2 |
| Region | chr16:3820941-3820941 | chr16 | 3820694 | 3820964 | p.G837E | CREBBP |
| Region | chr11:62398649-62398649 | chr11 | 62398401 | 62398671 | p.F238V | GANAB |
| Region | chr21:27141315-27141319 | chr21 | 27141079 | 27141344 | p.380 | GABPA |
| Region | chr8:145140619-145140619 | chr8 | 145140420 | 145140658 | p.A532V | GPAA1 |
| Region | chr20:23728500-23728500 | chr20 | 23728253 | 23728523 | p.W127R | CST1 |
| Region | chr12:13768566-13768566 | chr12 | 13768317 | 13768585 | p.K454T | GRIN2B |
| Region | chr14:96848786-96848786 | chr14 | 96848576 | 96848814 | p.V68L | GSKIP |
| Region | chr6:43018723-43018723 | chr6 | 43018473 | 43018743 | p.G490S | CUL7 |
| Region | chr12:104476535-104476535 | chr12 | 104476293 | 104476563 | p.K307KK | HCFC2 |
| Region | chr1:201180134-201180134 | chr1 | 201179920 | 201180185 | p.L2038* | IGFN1 |
| Region | chr20:10620342-10620342 | chr20 | 10620102 | 10620372 | p.S1154C | JAG1 |
| dbSNP | rs112710776 | chr1 | 64698052 | 64698278 | CNV | CNV |
| Region | chr6:43016239-43016239 | chr6 | 43015994 | 43016260 | p.G716W | CUL7 |
| Region | chr1:65301158-65301158 | chr1 | 65300914 | 65301180 | p.G1097D/V | JAK1 |
| Region | chr12:121891151-121891159 | chr12 | 121890915 | 121891180 | SpliceAccDon | KDM2B |
| Region | chr6:17809082-17809082 | chr6 | 17808832 | 17809102 | p.F694V | KIF13A |
| Region | chr6:39839319-39839319 | chr6 | 39839071 | 39839341 | p.D339N | DAAM2 |
| Region | chr7:151970885-151970885 | chr7 | 151970641 | 151970911 | p.Y306C | KMT2C |
| Region | chrX:41205658-41205658 | chrX | 41205420 | 41205690 | p.T498A | DDX3X |
| Region | chr12:49416133-49416133 | chr12 | 49415886 | 49416156 | p.R5448* | KMT2D |
| Region | chr12:52913543-52913543 | chr12 | 52913306 | 52913576 | p.A180S | KRT5 |
| Region | chr1:209791344-209791344 | chr1 | 209791108 | 209791365 | p.L987V | LAMB3 |
| Region | chr1:32751133-32751133 | chr1 | 32750906 | 32751154 | p.V479A | LCK |
| Region | chr19:14281521-14281521 | chr19 | 14281323 | 14281542 | p.E123K | LPHN1 |
| Region | chr20:39985707-39985707 | chr20 | 39985481 | 39985728 | p.N611D | LPIN3 |
| Region | chr12:57577605-57577605 | chr12 | 57577377 | 57577634 | p.R1948W | LRP1 |
| Region | chr2:170131572-170131572 | chr2 | 170131322 | 170131592 | p.V650X | LRP2 |
| Region | chrX:35821032-35821032 | chrX | 35820784 | 35821054 | p.H240P | MAGEB16 |
| Region | chr3:65415794-65415794 | chr3 | 65415547 | 65415817 | p.N523S | MAGI1 |
| Region | chr11:63662754-63662754 | chr11 | 63662506 | 63662776 | p.G27S | MARK2 |
| Region | chr9:36594637-36594637 | chr9 | 36594397 | 36594667 | p.G92R | MELK |
| Region | chr1:156051375-156051375 | chr1 | 156051135 | 156051405 | p.P139T | MEX3A |
| Region | chr1:222803356-222803356 | chr1 | 222803106 | 222803376 | p.E932* | MIA3 |
| Region | chr11:12278386-12278386 | chr11 | 12278155 | 12278417 | p.F814V | MICAL2 |
| Region | chr4:90857296-90857298 | chr4 | 90857057 | 90857327 | p.K822X | MMRN1 |
| Region | chr17:41958182-41958182 | chr17 | 41957958 | 41958217 | p.A360S | MPP2 |
| Region | chr3:121563345-121563345 | chr3 | 121563196 | 121563391 | p.L51R | EAF2 |
| Region | chr4:100510864-100510864 | chr4 | 100510619 | 100510889 | p.A153D | MTTP |
| Region | chr9:88628754-88628754 | chr9 | 88628510 | 88628780 | p.L504F | NAA35 |
| Region | chr8:24772021-24772021 | chr8 | 24771796 | 24772066 | p.E239K | NEFM |
| Region | chr11:20981980-20981980 | chr11 | 20981733 | 20982003 | p.H392N | NELL1 |
| Region | chr3:52519806-52519806 | chr3 | 52519576 | 52519826 | p.D556Y | NISCH |
| Region | chrX:105153557-105153557 | chrX | 105153310 | 105153580 | p.G642* | NRK |
| Region | chr9:23762076-23762076 | chr9 | 23761852 | 23762098 | p.E53* | ELAVL2 |
| Region | chr12:121618259-121618259 | chr12 | 121618073 | 121618343 | SpliceAccDon | P2RX7 |
| Region | chr9:118973971-118973971 | chr9 | 118973724 | 118973994 | p.M560V | PAPPA |
| Region | chr10:35928677-35928677 | chr10 | 35928566 | 35928756 | p.W561R | FZD8 |
| Region | chr7:106523507-106523507 | chr7 | 106523261 | 106523531 | p.T887A | PIK3CG |
| Region | chr19:18273259-18273259 | chr19 | 18273046 | 18273281 | p.T351I | PIK3R2 |
| Region | chrX:136112596-136112596 | chrX | 136112348 | 136112618 | p.G413V | GPR101 |
| Region | chr10:95931073-95931073 | chr10 | 95930825 | 95931095 | p.E235D | PLCE1 |
| Region | chr10:115341835-115341835 | chr10 | 115341696 | 115341966 | p.C347S | HABP2 |
| Region | chr1:65325832-65325834 | chr1 | 65325719 | 65325969 | p.PV430fs | JAK1 |
| Region | chr6:106547237-106547237 | chr6 | 106546987 | 106547257 | p.W24* | PRDM1 |
| Region | chr17:66519918-66519918 | chr17 | 66519692 | 66519962 | p.K134T | PRKAR1A |
| Region | chr14:30194762-30194762 | chr14 | 30194514 | 30194784 | p.L128R | PRKD1 |
| Region | chr2:128186178-128186178 | chr2 | 128185930 | 128186200 | p.R348* | PROC |
| Region | chr1:198687310-198687310 | chr1 | 198687098 | 198687333 | p.G513A | PTPRC |
| Region | chr9:8375948-8375950 | chr9 | 8375704 | 8375974 | p.V1550fs | PTPRD |
| Region | chr19:51480925-51480925 | chr19 | 51480706 | 51480966 | p.V210E | KLK7 |
| Region | chr12:49443667-49443668 | chr12 | 49443447 | 49443717 | p.G1235fs | KMT2D |
| Region | chr5:131939082-131939084 | chr5 | 131938861 | 131939123 | p.D628X | RAD50 |
| Region | chr2:173900884-173900884 | chr2 | 173900639 | 173900906 | p.L911P | RAPGEF4 |
| Region | chr16:816681-816681 | chr16 | 816432 | 816702 | p.I415T | MSLN |
| Region | chr10:12214837-12214837 | chr10 | 12214592 | 12214857 | p.N138K | NUDT5 |
| Region | chr5:140857741-140857750 | chr5 | 140857516 | 140857782 | p.K687X | PCDHGC3 |
| Region | chr1:237494245-237494245 | chr1 | 237494002 | 237494265 | p.Q79R | RYR2 |
| Region | chr14:81943431-81943434 | chr14 | 81943183 | 81943453 | p.T757NS | SEL1L |
| Region | chr12:122242657-122242666 | chr12 | 122242457 | 122242727 | p.H5HP | SETD1B |
| Region | chr11:94322285-94322285 | chr11 | 94322043 | 94322313 | SpliceAccDon | PIWIL4 |
| Region | chr11:65410070-65410070 | chr11 | 65409818 | 65410088 | p.P315L | SIPA1 |
| Region | chr11:16812425-16812425 | chr11 | 16812191 | 16812449 | p.A992E | PLEKHA7 |
| Region | chr14:23818598-23818598 | chr14 | 23818349 | 23818619 | p.L137M | SLC22A17 |
| Region | chr19:11134269-11134269 | chr19 | 11134030 | 11134291 | p.R979* | SMARCA4 |
| Region | chr7:127344940-127344969 | chr7 | 127344738 | 127345008 | p.VDWSIAVYT | SND1 |
| Region | chr17:76354701-76354722 | chr17 | 76354487 | 76354757 | p.SAQPLPG152X | SOCS3 |
| Region | chr1:167339423-167339423 | chr1 | 167339180 | 167339445 | p.Q85L | POU2F1 |
| Region | chr17:40474419-40474419 | chr17 | 40474197 | 40474448 | p.D661V | STAT3 |
| Region | chr17:40474420-40474420 | chr17 | 40474197 | 40474448 | p.D661Y | STAT3 |
| Region | chr17:40475063-40475063 | chr17 | 40474818 | 40475088 | p.E616G | STAT3 |
| Region | chr17:40369261-40369261 | chr17 | 40369015 | 40369285 | p.E433K | STAT5B |
| Region | chr10:96267065-96267065 | chr10 | 96266824 | 96267094 | p.K521E | TBC1D12 |
| Region | chr16:10788258-10788258 | chr16 | 10788013 | 10788282 | p.E158A | TEKT5 |
| dbSNP | rs112701466 | chr4 | 104510642 | 104510912 | CNV | CNV |
| dbSNP | rs112035841 | chr4 | 105412679 | 105412869 | CNV | CNV |
| dbSNP | chr4:106190722-106190923 | chr4 | 106190694 | 106190944 | CNV | TET2 |
| dbSNP | chr4:106180740-106180926 | chr4 | 106180717 | 106180954 | CNV | TET2 |
| dbSNP | chr4:106182804-106183005 | chr4 | 106182776 | 106183030 | CNV | TET2 |
| Region | chr4:106158284-106158287 | chr4 | 106158127 | 106158397 | p.VT1062Vfs | TET2 |
| Region | chr2:74274355-74274355 | chr2 | 74274107 | 74274377 | p.W302* | TET3 |
| Region | chr9:8341126-8341126 | chr9 | 8340879 | 8341149 | p.S1697Y | PTPRD |
| Region | chr7:11446625-11446625 | chr7 | 11446379 | 11446649 | p.C1325F | THSD7A |
| Region | chr1:226109259-226109259 | chr1 | 226109013 | 226109283 | p.A209G | PYCR2 |
| Region | chr12:94975755-94975755 | chr12 | 94975505 | 94975775 | p.K213T | TMCC3 |
| Region | chr4:159156626-159156626 | chr4 | 159156381 | 159156651 | p.D178H | TMEM144 |
| Region | chr4:3424263-3424263 | chr4 | 3424113 | 3424375 | p.A352V | RGS12 |
| Region | chr1:192627495-192627495 | chr1 | 192627278 | 192627519 | p.E98K | RGS13 |
| Region | chr3:158178791-158178791 | chr3 | 158178568 | 158178838 | p.E159D | RSRC1 |
| Region | chr12:111884606-111884606 | chr12 | 111884357 | 111884627 | p.R261Q | SH2B3 |
| Region | chr20:44681605-44681605 | chr20 | 44681375 | 44681626 | p.R819L | SLC12A5 |
| Region | chr4:167675857-167675857 | chr4 | 167675619 | 167675877 | p.I245F | SPOCK3 |
| Region | chr6:138192466-138192471 | chr6 | 138192228 | 138192498 | p.G35GG | TNFAIP3 |
| Region | chrX:122745295-122745295 | chrX | 122745049 | 122745319 | p.K1584Q | THOC2 |
| Region | chr6:138198315-138198322 | chr6 | 138198155 | 138198405 | p.L303LK | TNFAIP3 |
| Region | chr16:19452100-19452100 | chr16 | 19451859 | 19452125 | p.E247G | TMC5 |
| Region | chr6:138192466-138192472 | chr6 | 138192228 | 138192498 | p.GG35fs | TNFAIP3 |
| Region | chr4:122068326-122068326 | chr4 | 122068101 | 122068355 | p.Q275* | TNIP3 |
| Region | chr4:122078287-122078287 | chr4 | 122078067 | 122078337 | p.Q179* | TNIP3 |
| Region | chr3:195609046-195609046 | chr3 | 195608796 | 195609066 | p.A255T | TNK2 |
| Region | chr5:114462264-114462264 | chr5 | 114462016 | 114462286 | p.S708N | TRIM36 |
| Region | chr4:189012946-189012946 | chr4 | 189012714 | 189012978 | p.F249V | TRIML2 |
| Region | chr13:38211101-38211101 | chr13 | 38210868 | 38211129 | p.D785G | TRPC4 |
| Region | chr16:2112528-2112528 | chr16 | 2112307 | 2112568 | p.R430* | TSC2 |
| Region | chr21:38537997-38537997 | chr21 | 38537759 | 38538029 | p.R1161C | TTC3 |
| Region | chr2:179427933-179427933 | chr2 | 179427692 | 179427962 | p.Y18769* | TTN |
| Region | chr3:126366093-126366093 | chr3 | 126365846 | 126366116 | p.E134G | TXNRD3 |
| Region | chr8:103291364-103291364 | chr8 | 103291141 | 103291392 | p.R2025H | UBR5 |
| Region | chr11:18579781-18579781 | chr11 | 18579536 | 18579806 | p.S215G | UEVLD |
| Region | chr4:85617226-85617226 | chr4 | 85616981 | 85617251 | p.H2933P | WDFY3 |
| Region | chr18:54424496-54424496 | chr18 | 54424253 | 54424523 | p.L891P | WDR7 |
| Region | chr16:72991453-72991453 | chr16 | 72991216 | 72991473 | p.Y864* | ZFHX3 |
| Region | chr19:22155948-22155948 | chr19 | 22155704 | 22155974 | p.I630F | ZNF208 |
| Region | chr20:52198240-52198240 | chr20 | 52197990 | 52198260 | p.H376D | ZNF217 |
| Region | chr19:53269181-53269181 | chr19 | 53268944 | 53269212 | p.Y610H | ZNF600 |
| Region | chr19:53995049-53995049 | chr19 | 53994838 | 53995107 | p.K521N | ZNF813 |
| Region | chr12:133634520-133634520 | chr12 | 133634276 | 133634546 | p.E407* | ZNF84 |
| Region | chr16:3139621-3139621 | chr16 | 3139573 | 3139763 | p.P468L | ZSCAN10 |

### Supplementary Table 2: Single-cell DNA sequencing custom panel amplicon coverage
