## Supplemental MATERIAL AND METHODS for "CLONAL LYMPHOCYTE EXPANSIONS AND JAK-STAT PATHWAY MUTATIONS DEFINE A PATHOGENIC CONTINUUM DRIVING RESISTANCE TO GLUTEN-FREE DIET IN CELIAC DISEASE"

#### Patient enrolment and tissue sampling

Blood and duodenal biopsies were collected during diagnostic and follow-up endoscopic procedures at AP-HP Cochin Hospital according to the ENTEROPATH protocol Inserm C19-07 (IDRCB :2019-A01594-53; N°CPP :2019/56). Diagnosis of CeD was based on positive anti-transglutaminase IgA serology, and histological criteria of Marsh classification at initial diagnosis[1,2]. All tested patients carried the HLA-DQ2 or -DQ8 haplotype (4/22). Patient characteristics are presented in Table 1. Active CeD was newly diagnosed in a patient consulting in our referral tertiary center. Diagnosis of RCD was made in CeD patients with persistent symptoms and villous atrophy despite strict GFD for more than one year and classified by flow cytometry as RCD1 if >75% of CD103+ IELs were $CD8^{+}sCD3^{+}TCR\alpha\beta^{+}$ and RCD2 when over 30% $CD103^{+}$ IELs contained intracellular CD3ε but lacked surface CD3-TCR complexes (11 cases) or, in one atypical case when >90% IELs expressed a TCRγδ. Duodenal biopsies contained clonal TCRG rearrangements in all RCD2 patients and oligoclonal TCRG rearrangements in 1/4 RCD1. Controls were individuals with normal duodenal histology in whom CeD and other autoimmune/inflammatory disease were excluded.

#### Sample processing

Peripheral blood was collected in heparin-coated BD Vacutainer tubes, and PBMCs were isolated by Ficoll gradient centrifugation, then cryopreserved in CryoStor® CS10 at −80°C and stored in liquid nitrogen. Duodenal biopsies (5–6 per patient) were placed directly into CryoStor® CS10, snap-frozen, and stored at −80°C prior to processing.

#### Cell Type Annotation and Surface Protein Analysis

Cell type labels were assigned to clusters based on a manually curated list of marker genes and known immune and tumor cell signatures from published data[3]. A total of 217,267 cells were retained for further analysis, including 45,248 tumor cells and 172,019 non-tumor cells. To evaluate the expression of 21 to 130 surface proteins (TotalSeq™-C Human Universal Cocktail, V1.0, Biolegend), antibody-derived tag (ADT) counts were incorporated into the integrated Seurat object and normalized using the centered log-ratio (CLR) transformation method.

**Differential expression analysis**

Differential expression analysis was conducted on log-normalized, non-batch-corrected data using the Wilcoxon rank-sum test with Bonferroni correction, as implemented in the *FindMarkers* function. Genes with adjusted p-values < 0.05 were considered significant. Differentially expressed genes (DEGs) were further classified as upregulated or downregulated based on the log2 fold change (avg_log2FC > 0 for upregulated genes and avg_log2FC < 0 for downregulated genes). Signature scores were computed using the *AddModuleScore_UCell* function from the UCell package. All figures were generated using a combination of Scpubr (v2.0.2) and ggplot2.

**InferCNV analysis**

*InferCNV* was used to identify large-scale somatic chromosomal copy number variations (CNVs)[4]. An *infercnv* object was created by combining the raw counts extracted from the integrated Seurat object (RNA assay and counts slot) with the positional information of genes from the human genome reference (hg38_gencode_v27). Healthy cells were used as the reference group, and CNVs were inferred using the 6-state hidden Markov model (i6). Sex chromosomes (chrX and chrY) and mitochondrial DNA (chrM) were excluded from the analysis.

### RNA velocity and trajectory reconstruction

BAM files for each sample were processed using the *Velocyto* software[5] with default parameters to compute RNA velocity. The resulting loom files were integrated and analyzed with *scVelo*[6]. The average velocity values, calculated using the dynamical model, were projected onto UMAP embeddings. Cells were ordered by latent time, and root cells were selected from the velocity embedding grid. Pseudotime analysis was then performed using *Monocle3*[7,8].

### Cell sorting for exome sequencing

Intestinal cells and PBMCs from patients P10 and P43 were stained with the cocktail Vβ antibody F (containing the anti-human Vβ1-FITC/PE, clone BL37.2, for P10) or E (containing the anti-human Vβ2-FITC/PE, clone MPB2D5, for P43) from the Beta Mark TCR Vbeta Repertoire Kit (Beckman Coulter) on ice for 20 minutes. After one wash in PBS added with 2% human AB serum, cells were stained with anti-human CD45-PerCP/Cy5.5 (clone HI30), anti-human CD3-BV650 (clone SK7), anti-human CD103-BV605 (clone Ber-ACT8), anti-human CD4-PE/Cy7 (clone RPA-T4), all from SONY Biotechnology Inc, anti-human CD8-APC/H7 (clone SK1, BD Biosciences) and fixable aqua LIVE/DEAD (ThermoFisher Scientific). For P10, live $CD45^+CD3^+CD8^+V\beta1^+$, $CD45^+CD3^+CD8^+V\beta1^-$ and $CD45^+CD3^+CD4^+$ $V\beta1^-$ cells were sorted from PBMCs and duodenal biopsy-derived cells using the Cell SONY MA900 Multi-Application Cell Sorter. For P43, live $CD45^+sCD3^-CD103^+$ RCD2 cells and $CD45^+CD3^+CD4^+V\beta2^-$ (control cells) were sorted from isolated duodenal cells and $CD45^+sCD3^-CD103^+$ RCD2 cells, $CD45^+CD3^+CD4^+V\beta2^+$ cells ($CD4^+$ lymphoproliferation), and $CD45^+CD3^+CD4^+V\beta2^-$ and $CD45^+CD3^+CD8^+V\beta2^-$ (control cells) were sorted from PBMCs. Dried pellets were kept at -20°C for exome sequencing.

**Single-Cell DNA Sequencing**

ScDNA-seq was performed using the Mission Bio Tapestri scDNA-seq V2 platform, following the manufacturer's instructions. A custom ScDNA-seq panel targeting 210 mutations and chromosomal structural abnormalities identified in RCD2 patients[9] (**Supplementary Table 2**) was designed with and manufactured by Mission Bio, Inc. Approximately 120,000 $CD45^{+}sCD3^{-}CD103^{+}$ RCD2 sorted from thawed PBMCs of one patient with 80% circulating RCD2 cells (P4 gut) and 50,000 cells from tumor cell lines derived from the blood and biopsy samples of the same patient were resuspended in cell buffer and used for microfluidic encapsulation, lysis, and cell barcoding on the Tapestri platform. Targeted DNA regions were amplified by incubating the barcoded DNA emulsions in a thermocycler with the following program: 98°C for 6 minutes (4°C/sec); 11 cycles of 95°C for 30 seconds, 72°C for 10 seconds, 61°C for 3 minutes, 72°C for 20 seconds (1°C/sec); 13 cycles of 95°C for 30 seconds, 72°C for 10 seconds, 48°C for 3 minutes, 72°C for 20 seconds (1°C/sec); and 72°C for 6 minutes (4°C/sec). Emulsions were broken, and DNA was digested and purified with 0.42X Ampure XP reagent (Beckman Coulter). The beads were pelleted, washed with 80% ethanol, and DNA targets were eluted in nuclease-free water. Indexed Illumina libraries were generated by amplifying DNA libraries with Mission Bio V2 Index Primers in a thermocycler using the following program: 95°C for 3 minutes; 10 cycles (DNA library) or 20 cycles (protein library) of 98°C for 20 seconds, 62°C for 20 seconds, 72°C for 45 seconds; and 72°C for 2 minutes. Final libraries were purified with 0.42X Ampure XP reagent. Libraries were pooled and subjected to paired-end 150-bp sequencing on a Novaseq 6000 (Illumina). Raw FASTQ files were analyzed using the Tapestri pipeline (Mission Bio) and custom Python scripts.

**Generation of ruxolitinib-resistant RCD2 cell lines**

The intestinal RCD2 cell line analyzed by single cell genomics  was cultured in RPMI supplemented with 10% human AB serum (Sigma Aldrich), 1% sodium pyruvate, 1% non-essential amino acids, 1% HEPES buffer, 1 µg/mL fungizone, 40 µg/mL gentamicin, 5x$10^5$ M β-mercaptoethanol (Invitrogen), and 20 ng/mL human IL-15 (R&D Systems). Ruxolitinib-resistant cell lines were generated through three cycles of treatment with 400 nM ruxolitinib (Selleckchem, Euromedex, France). Each treatment cycle lasted one week, during which cells were first resuspended in the presence of ruxolitinib in 24-well plates and wells completed with fresh medium without ruxolinitib every 2-3 days. Each cycle was followed by a three-week recovery period in medium without ruxolitinib. Control cell lines were obtained in parallel cultures where ruxolitinib was replaced by the vehicle DMSO (Sigma-Aldrich, USA).

Apoptosis was determined using AnnexinV-APC and propidium iodide (Apoptosis detection kit, SONY Biotechnology Inc) after 24H and 48H of treatment with 400nM ruxolitinib or DMSO. Data were acquired on a FACSCantoII (BD Biosciences) flow cytometer and analysed with FlowJo version 10 software.

**Statistics**

Statistical analyses were performed using Prism version 6 (GraphPad Software, La Jolla, CA, USA). Multiple group comparisons were conducted using the Kruskal–Wallis test followed by Dunn’s post hoc test for multiple comparisons. A corrected $p$ value $< 0.05$ was considered statistically significant.

**References**

1. Marsh MN. Gluten, major histocompatibility complex, and the small intestine. A molecular and immunobiologic approach to the spectrum of gluten sensitivity ('celiac

sprue'). Gastroenterology. 1992;102(1):330-354.
