## Supplementary Table Legends for "CLONAL LYMPHOCYTE EXPANSIONS AND JAK-STAT PATHWAY MUTATIONS DEFINE A PATHOGENIC CONTINUUM DRIVING RESISTANCE TO GLUTEN-FREE DIET IN CELIAC DISEASE"

**Supplementary Table S3. Somatic mutations enriched in RCD2 cell lines following ruxolitinib treatment.**

**Supplementary Table S4.**

**Supplementary Table S5. Somatic variants identified in the Vβ2$^+$ CD4$^+$ lymphoproliferation clone from patient P43.**

Whole-exome sequencing of sorted Vβ2$^+$ CD4$^+$ T cells revealed 41 de novo somatic mutations relative to Vβ2$^-$ control T cells. Notably, this included a heterozygous missense mutation in the tumor suppressor gene *ATM* (c.7422A>C, p.L2474F), predicted to be deleterious with a CADD score of 25. The presence of mutations in genes such as *ATM*, which is frequently altered in lymphoid malignancies, supports the hypothesis that this population represents a transformed lymphocyte clone.

**Supplementary Table S6. Somatic mutations in the SOCS1 and SOCS3 genes identified in the expanded intestinal Vβ1$^+$ CD8$^+$ T-cell clone from a patient with RCD1.**

Whole-exome sequencing of sorted Vβ1$^+$ CD8$^+$ T cells from duodenal and blood samples revealed two somatic mutations absent from Vβ1$^-$ CD8$^+$ control T cells: a frameshift mutation in *SOCS1* (c.129delC, p.P43fs) and a nonsense mutation in *SOCS3* (c.259C>T, p.Q87*). Both genes are key negative regulators of the JAK-STAT pathway, and their inactivation is predicted to enhance cytokine signaling in the inflamed gut environment, potentially driving clonal expansion, cytotoxic activity, and sustained mucosal damage in RCD1.
